# Site-specialization of human oral *Porphyromonas* species

**DOI:** 10.64898/2026.06.02.729646

**Authors:** Julian Torres-Morales, Floyd E. Dewhirst, Kathryn M. Kauffman, Jessica L. Mark Welch, Gary G. Borisy

## Abstract

Site-specificity within the human oral cavity reflects adaptation mechanisms such as genome divergence and metabolic specialization. Members of the genus *Porphyromonas* are distributed across oral sites in health and disease, yet the specific distribution of taxa and the functional basis of their site-specificity remain poorly understood. We analyzed 1,242 metagenomes from nine oral sites in healthy individuals and 24 subgingival plaque samples from individuals with periodontitis. Competitive mapping to a dereplicated genus-level pangenome of 84 reference genomes, combined with phylogenomic, gene-level detection, and functional profiling, revealed distinct site-specific distribution patterns, ecotype differentiation, and metabolic specialization across *Porphyromonas* taxa. *Porphyromonas pasteri* was the most abundant and widespread taxon in healthy subjects, comprising two ecotypes--one mucosal, one plaque-associated. *Porphyromonas gingivalis* was rare in healthy subjects but present in periodontal disease, although detected in only half of periodontitis samples. *P*. *gingivalis* exhibited the broadest metabolic repertoire, suggestive of a survival strategy adaptive to disparate conditions. In contrast, *Porphyromonas catoniae*, restricted to healthy dental plaque, lacked biosynthetic pathways for cobalamin, biotin, and serine, implying nutritional dependency on other taxa or the host. *Porphyromonas endodontalis*, detected in subgingival plaque across both health and disease, also lacked several metabolic pathways. A 44 kb conjugative element identified in *P*. *gingivalis* was detected across healthy and periodontitis subgingival plaque microbiomes independently of the *P*. *gingivalis* chromosome, indicating horizontal transfer. These findings reveal genomic divergence and complex metabolic specialization among *Porphyromonas* taxa, refining our understanding of their role in the ecological structure of the human oral microbiome.

**Importance:** Microbial specialization within host-associated ecosystems is a key driver of community structure and function. The human oral cavity, with its spatially structured and ecologically diverse niches, offers a powerful model for studying these patterns. Our findings reveal that *Porphyromonas* taxa are largely restricted to specific oral sites, with *Porphyromonas pasteri* emerging as the most abundant and ecologically versatile member of the genus, and that large mobile elements shape genomic diversity across oral taxa. This work advances our understanding of microbial biogeography and highlights the ecological and metabolic significance of *Porphyromonas* taxa within the complex oral community.

## Introduction

The human oral microbiome is spatially organized across anatomical and physico-chemically distinct habitats–teeth, tongue, gingival sulcus, and mucosal surfaces–each imposing selective pressures that contribute to the assembly of distinctive communities at each oral site (1–3). Within the mouth, closely related taxa specialize to distinct oral niches; for example, one species will localize primarily to subgingival and supragingival dental plaque, and a closely related species will localize to mucosal sites such as the tongue, tonsils, and throat. This pattern, formalized as the site-specialist hypothesis (4), can be recognized when the data are analyzed with sufficient genomic resolution. Evidence from cultivation-based surveys, imaging, and metagenomic analyses supports this framework for several oral genera, linking genomic variation within closely related species to biogeographic distributions within the mouth (5–13).

Members of the genus *Porphyromonas*, including *Porphyromonas gingivalis*, *Porphyromonas endodontalis*, *Porphyromonas catoniae*, and *Porphyromonas pasteri*, are key taxa of the human oral microbiome in health and disease (2, 14–18). Although much effort in oral microbiology has been focused on *P. gingivalis* and *P. endodontalis* because of their association with periodontal disease, they are rare in healthy individuals (19). In contrast, *P. pasteri* is far more prevalent and abundant in the healthy oral microbiome and is broadly distributed across multiple oral sites (4, 20–22), raising the question of whether *P. pasteri* is an exception to the site-specialist hypothesis or reflects unresolved subpopulation structure (23). Phylogenetic analysis of ancient and modern metagenome-assembled genomes (MAGs) has suggested tropism of *P. pasteri* clades to tongue or plaque; however, host-level ecological distributions of these subpopulations *in situ* cannot be resolved from phylogenies alone (24).

As a foundation for addressing the role of *Porphyromonas* in the oral microbiome, we investigated the distribution, abundance, and functional capacity of members of the genus, including its highly abundant and understudied members, in both health and disease. We carried out a systematic analysis using metapangenomics to link population genomic variation to ecological distributions across the genus. By combining pangenome analysis with competitive metagenomic read recruitment, metapangenomics enables simultaneous identification of habitat-specific populations, detection of ecologically relevant gene content, and characterization of genomic features, including mobile genetic elements, associated with environmental adaptation. This approach has revealed ecotype differentiation and habitat-specific gene content in both marine microbes and several oral taxa (7–13, 25, 26). Here, we apply the metapangenomic framework across 1,242 human oral metagenomes spanning nine healthy oral sites and periodontitis-associated subgingival plaque to systematically resolve ecological differentiation across *Porphyromonas*. We show that species-level niche partitioning is associated with distinct metabolic strategies; that *P. pasteri* comprises two ecologically distinct ecotypes despite minimal gene-content and functional variation; and that a discrete ∼44 kb mobile genomic island, first detected in the *P*. *gingivalis* genome (27, 28), encodes the full machinery for autonomous transfer and persists in healthy subgingival plaque across genera, independently of the *P*. *gingivalis* genome.

## Results

### A curated pangenomic reference for human oral *Porphyromonas*

To resolve ecological differences among human oral *Porphyromonas* taxa, we downloaded and evaluated 377 publicly available *Porphyromonas* genomes including those of both human and non-human origin, retaining 84 reference genomes associated with the human oral cavity after quality filtering, phylogenomic placement, taxonomic assignment, and 98% average nucleotide identity (ANI) dereplication (see Methods, Extended Data, and Tables S1-S9). The evolutionary and taxonomic framework underlying this reference set is described in detail in the Extended Data, including a phylogeny of the 343 *Porphyromonas* genomes after quality filtering (Extended Data Fig. E1). Ecological characterization of type strains of human origin, including non-oral *Porphyromonas*, is shown based on mapping of metagenomic reads (Extended Data Fig. E2). After eliminating genomes of non-human origin, a pangenome of 191 genomes, prior to dereplication, of *Porphyromonas* strains associated with the human oral cavity is also presented (Extended Data Fig. E3). After dereplication, 84 reference genomes associated with the human oral cavity were retained, and their predicted genes were grouped into homologous gene clusters based on amino acid sequence similarity, enabling systematic evaluation of gene-content variation across oral taxa (Fig. 1).

**Fig. 1.**
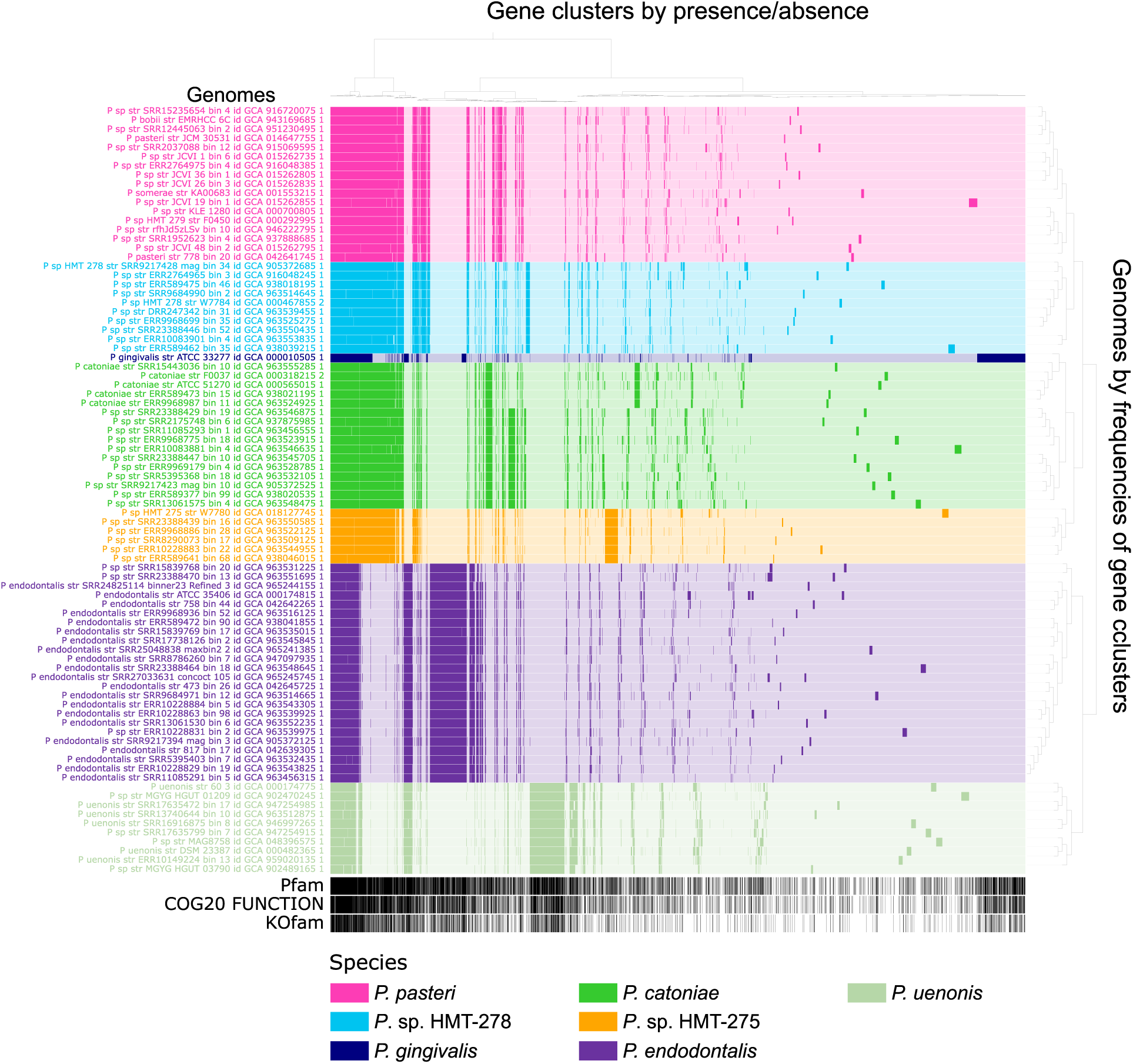
Dereplicated pangenome of human oral *Porphyromonas*. Pangenome analysis of 84 high-quality reference genomes obtained after dereplication at 98% ANI reveals well-defined genomic groups that correspond to species-level designations. Genes were predicted and translated with Prodigal, compared using BLASTp, and clustered into homologous gene clusters using a Markov Clustering Algorithm (MCL) with an inflation parameter of 10. Gene cluster presence (solid colors) and absence (pale segments) are shown for each genome as vertical bars. Genomes are color-coded by genomic group; both genomes and gene clusters are hierarchically clustered by gene cluster frequency and presence/absence patterns, respectively. Functional annotations from Pfam, NCBI COG, and KEGG KOfam are indicated by black bars below each gene cluster.

The pangenome shows which gene clusters are core, present in all genomes, or accessory, present in only a subset. Hierarchical clustering of genomes based on these presence-absence patterns revealed coherent genomic groups, most corresponding to recognized species: *P. pasteri*, *P*. *catoniae*, *P*. *gingivalis*, *P*. *endodontalis*, and *P*. *uenonis*; as well as human microbial taxon (HMT) group designations: *P*. sp. HMT-275 and *P*. sp. HMT-278. *P. catoniae,* however, was divisible into two distinguishable pangenomic groups.

We resolved several misclassifications of genomes based on the pangenome clustering. Within *P. pasteri* (n = 17 genomes), only two were originally deposited under that name, while twelve unnamed *Porphyromonas* sp. genomes clustered within this taxon. In addition, *P*. *somerae* strain KA00683 was found to be misclassified, and *P*. *bobii,* initially isolated from prostate secretion fluid (29), also grouped within *P. pasteri,* indicating that *P. bobii* is a later, heterotypic synonym for *P. pasteri*. *Porphyromonas* sp. oral taxon 279 F0450 was renamed to *P. pasteri* (18) but retains the name *P.* sp. at NCBI. Thus, *P. pasteri* was expanded and refined from two to seventeen genomes even after 98% ANI dereplication. In contrast, the 99 *P*. *gingivalis* genomes before dereplication collapsed to a single representative after dereplication, indicating low genome diversity (high sequence similarity) in the *P. gingivalis* dataset. High alignment coverage was observed across all 4,851 pairwise comparisons of the 99 *P. gingivalis* genomes (mean 85.7% ± 3.1%) confirming that high ANI values reflected broad, genome-wide sequence similarity rather than being restricted to a few conserved regions, and explaining why all 99 genomes collapsed to a single representative at 98% ANI dereplication. High-quality MAGs substantially broadened the genomic collection; for example, *P*. sp. HMT-278 was represented by only a single isolate, but eight MAGs, for a total of nine genomes.

Independent metrics, using the full 191 pre-dereplicated genomic set for broadest group support, confirmed the pangenome-based genomic groups. Within each of the genomic groups, genomes shared ANI of >= 95% (Fig. S1A and Table S10). An exception was the *P. catoniae* set of genomes, which formed three closely related genomic groups with inter-group ANI values between 90% and 95%, indicating relatedness at a nucleotide level below the classic species boundary. A tanglegram comparing pangenome-based clustering to a phylogeny based on 61 single-copy core genes (Fig. S1B) revealed concordance between phylogeny and gene content relationships, supporting the pangenomic grouping. Their ecological distribution patterns, described below, provide additional support for these groups. Overall, gene-content clustering, ANI, and phylogenomic analyses were mutually consistent, establishing a robust framework for ecological interpretations.

### Ecological niche partitioning across *Porphyromonas* genomic groups

To assess habitat associations for each genomic group, we competitively mapped quality-filtered metagenomic reads from 1,242 samples (Table S11) spanning nine healthy human oral sites and periodontitis-associated subgingival plaque onto the 84 dereplicated reference genomes (see Methods). We quantified genome presence using a breadth of coverage metric. This metric measures the extent to which a reference genome matches natural populations in the metagenomes. Relative abundance of genomes was quantified using the mean depth of coverage of the nucleotides in the second and third quartiles of coverage (Q2-Q3) to minimize bias from cross-mapping to conserved and repetitive regions.

Distinct habitat associations were evident at the level of breadth of coverage (Fig. 2 and Table S12). *Porphyromonas pasteri* genomes displayed high breadth of coverage across all healthy sites, most frequently on the tongue dorsum, palatine tonsils, saliva, throat, and hard palate. In contrast, *P*. *gingivalis* showed high breadth of coverage in periodontitis samples and was found only rarely in healthy sites. *P*. *catoniae* was primarily associated with healthy sub-and supragingival plaque. *Porphyromonas endodontalis* was found in healthy sub- and supragingival plaque and periodontitis. *Porphyromonas* sp. HMT-275 and *P. uenonis* were rare at all sites. Applying a breadth threshold (≥ 25%) for genome detection (Fig. S2) did not change the qualitative conclusions. *Porphyromonas pasteri* was broadly detected across healthy sites, while *P*. *gingivalis* was present in only 10 of 24 (42%) periodontitis samples. *Porphyromonas catoniae* was detected mainly in healthy dental plaque, while *P. endodontalis* was detected mainly in supra- and sub-gingival plaque, tonsils, and in periodontitis.

**Fig. 2.**
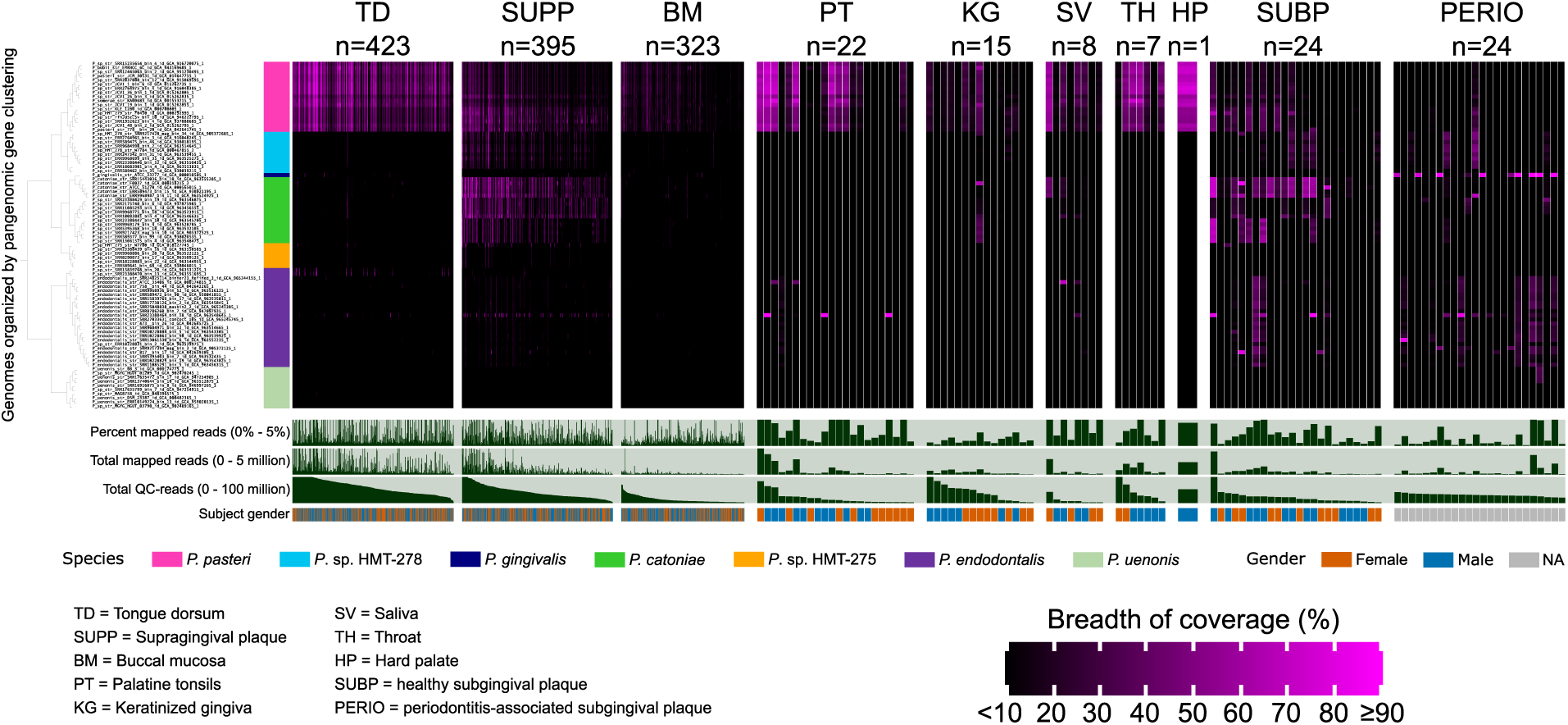
Breadth of coverage heatmap reveals site-specific distribution of *Porphyromonas* genomes. Competitive mapping of 1,242 human metagenomes from nine healthy oral sites and periodontitis-associated subgingival plaque to 84 dereplicated reference genomes reveals distinct ecological partitioning. The heatmap shows the fraction of genome nucleotides covered by at least one sequencing read, shown as a gradient from <10% (black) to ≥90% (magenta). Genomes are ordered by gene cluster frequency profiles (pangenome in Fig. 1); samples are ordered by decreasing total quality-filtered read count within each site.

The relative abundance of genomes within the *Porphyromonas* genus was determined from the mean depth of coverage (Fig S3 and Table S13). Aggregating per-genome relative abundance into genomic groups reduced within-group noise and revealed clear patterns of habitat partitioning, co-occurrence, and dominance (Fig. 3 and Tables S14, S15). *Porphyromonas pasteri* was broadly abundant across the healthy oral cavity but was divisible into two groups, one favoring mucosal sites, especially the tongue dorsum, and the other favoring dental plaque.

**Fig. 3.**
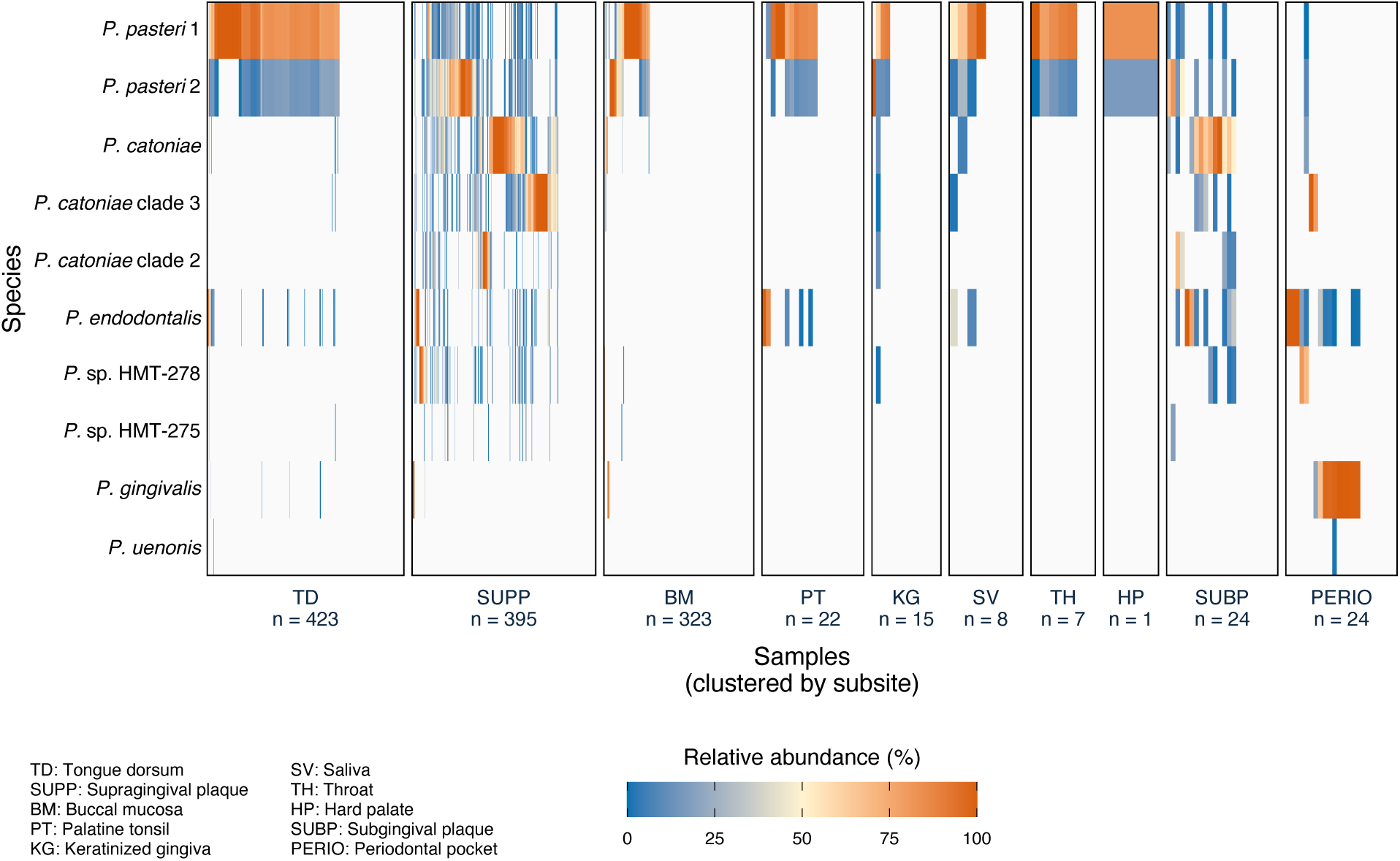
Species-level relative abundance reveals niche partitioning of *Porphyromonas* in the human oral cavity. Relative abundance was calculated as the sum of Q2-Q3 mean depth of coverage values for all genomes within each genomic group divided by the total Q2-Q3 depth of coverage for all *Porphyromonas* genomes (see Fig. S3 for genome-level data). Hierarchical clustering of samples within each site based on Bray-Curtis similarity reveals that genomic groups exhibit mutual exclusion across samples.

*Porphyromonas catoniae* and its sister clades were specialized to healthy dental plaque. *Porphyromonas endodontalis* was primarily a subgingival plaque specialist that reached high abundance in periodontitis. *Porphyromonas gingivalis* was rare in healthy individuals and was abundant in only a subset of periodontitis samples. Tongue dorsum and other mucosal sites were typically dominated by *P. pasteri* alone, whereas multiple taxa were frequent in dental plaque.

Within healthy supra- and subgingival plaque, as visualized by the heatmap of Figure 3, individual samples tended to be dominated by one taxon. To quantify these patterns, we applied Fisher’s exact test, correlation analyses, and abundance comparisons to the most abundant and prevalent healthy plaque specialists (Table S16). Presence/absence analyses showed that the dominant plaque-associated taxa frequently co-occurred within healthy supragingival plaque, with little evidence for strict mutual exclusion. However, strong negative abundance correlations were observed among several taxa (Spearman rho = −0.34 to −0.64; FDR < 4×10^−8^), indicating that while taxa could colonize the same habitat, individual samples were typically dominated by one taxon over another. The strongest co-occurrence pattern involved the two *P*. *pasteri* subgroups, which frequently co-occurred in supragingival plaque (OR = 62.65; FDR = 1.4 × 10^−39^) and tongue dorsum (OR = 20.5, p = 0.003), but displayed strong abundance asymmetry that differed by habitat: *P*. *pasteri* 2 was dominant in supragingival plaque while *P*. *pasteri* 1 dominated on the tongue dorsum (Wilcox p = 8.16×10^−20^). *Porphyromonas catoniae* showed the strongest negative abundance correlations with both *P*. *pasteri* subgroups (rho = −0.59 to −0.64; FDR <4×10^−24^) and outcompeted *P*. *pasteri* 1 in abundance when co-occurring in plaque (Wilcox FDR = 6.2×10^−8^). In periodontal disease samples, *P*. *gingivalis*, *P*. *endodontalis*, *P*. sp. HMT-278, and *P*. *catoniae* clade 2 exhibited a winner-takes-all pattern–with largely non-overlapping occurrence across samples, consistent with competitive exclusion and fine-scale niche partitioning. Notably, *P*. *gingivalis* and *P*. *endodontalis* were abundant in only a fraction of periodontitis samples– *P*. *gingivalis* in 10/24 and *P*. *endodontalis* in 7/24 samples–with *P*. *endodontalis* also present in healthy subgingival plaque samples (8/24 samples), reinforcing the view that periodontal disease reflects polymicrobial changes rather than the presence of a single microbe (30, 31).

### Metabolic divergence reflects site specialization

Habitat partitioning patterns reflect the metabolic potential of genomic groups. Across the genus, 43 complete KEGG metabolic modules (containing ≥ 75% of required enzymes) were identified, and metabolic profiles covaried with ecological distributions (Fig. S4 and Table S17).

The dental plaque specialist, *P*. *catoniae*, possessed the most restricted metabolic capacity (32 complete modules), lacking biosynthetic pathways for cobalamin, biotin, serine, histidine, and the shikimate pathway–indicating dependence on exogenous nutrients and likely cross-feeding from co-occurring taxa such as *Corynebacterium* or *Streptococcus* that dominate dental plaque (11). *Porphyromonas endodontalis* also displayed a restricted metabolic profile, lacking galactose and glycogen utilization, lipid A biosynthesis, and the energy production phosphate acetyltransferase-acetate kinase pathway (M00579), compensating for it with the production of acetyl-CoA from pyruvate (M00169) as an alternative energy route. These restrictions and the resulting metabolic dependence are consistent with an ecological strategy in which highly specialized co-occurring taxa have co-evolved, reducing their metabolic burden by relying on other members of the community. *Porphyromonas gingivalis* possessed the broadest metabolic profile, including degradation of galactose and glycogen, biosynthesis of cobalamin, biotin, and the shikimate pathway, along with lipid A biosynthesis and the most extensive mobilome among *Porphyromonas* (>100 mobile element-associated genes). This metabolic flexibility is consistent with genomic adaptation enabling persistence under fluctuating nutrient and immune conditions of disease-associated subgingival conditions and may underlie its capacity to colonize independently of a stable community.

### *Porphyromonas pasteri* comprises two ecologically distinct ecotypes

*P. pasteri* is the most prevalent and abundant *Porphyromonas* species across the healthy human oral cavity. Notably, read mapping revealed two subgroups with distinct ecological signatures: one predominantly abundant on the tongue dorsum and other mucosal sites (*P. pasteri* 1), and one enriched in healthy dental plaque (*P. pasteri* 2) (Fig. 3, Fig. S3, and Table S15). To further test the validity of this sub-species structure, we expanded the *P. pasteri* genomic dataset with seven quality-filtered dereplicated non-RefSeq genomes retrieved from GenBank (see Methods and Tables S18, S19). Increasing genomic sampling in each subpopulation improves phylogenetic support for clade boundaries and enhances ecological signal captured by competitive mapping, enabling a more robust distinction between inter-species subgroups with overlapping distributions.

Competitive mapping to this expanded set confirmed distinct ecological preferences for each subgroup (Fig. 4A). *P. pasteri* 1 (13 genomes) was consistently abundant on the tongue dorsum and rarely detected in dental plaque. Meanwhile, *P. pasteri* 2 (11 genomes) was enriched in supragingival plaque and detected at a lower relative abundance on the tongue dorsum. This sub-species structure corresponds to two phylogenetically distinct subpopulations separated into two clades by phylogenomic analysis of 235 single-copy core genes (Fig. 4B). Intra- and inter-clade pairwise ANI indicated genomic divergence sufficient to support distinct ecological and evolutionary identities (Fig. 4C).

**Fig. 4.**
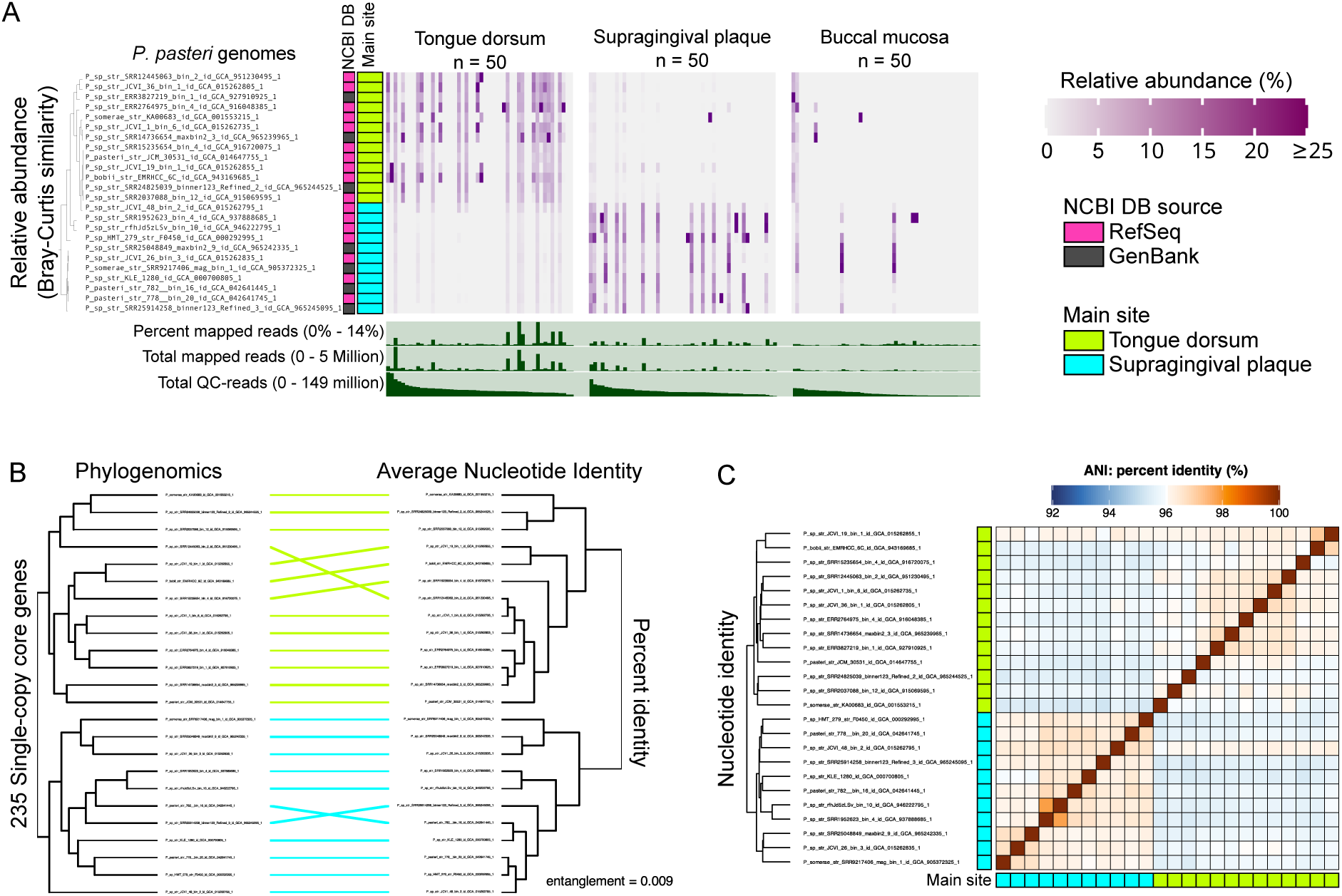
Genomic and ecological resolution of *P. pasteri* ecotypes. (A) Relative abundance of individual genomes across tongue dorsum (TD), supragingival plaque (SUPP), and buccal mucosa (BM) confirms site-specific ecological preferences. The 50 metagenomes with the highest *Porphyromonas* mapping read counts were selected per site. Genomes are clustered by distribution patterns using Euclidean distance and Ward linkage. (B) Tanglegram comparing phylogenomic and ANI-based clustering reveals congruence between both classification approaches. The phylogenomic tree was constructed from single-copy core genes (n=235) using the Whelan and Goldman substitution model with 1,000 bootstrap replicates (UFBoost). (C) ANI heatmap shows high intra-clade similarity and inter-clade divergence, with genomes ordered symmetrically along both axes.

Despite the differing site preferences of the two *P. pasteri* clades, neither gene-content clustering nor metabolic capacity distinguished the two subpopulations (Fig. S5, S6). Both lack biotin synthesis, histidine degradation, and the shikimate pathway, and both carry overlapping complements of mobile elements (n = 15 mobile-element genes per clade). Functional enrichment analysis across all genomic annotations–COG, Pfam, KOfam, CAZyme, and KEGG modules–identified a single function significantly enriched in *P. pasteri* 2: leucine carboxyl methyltransferase, present in all genomes from this subgroup and absent in *P. pasteri* 1 (Tables S20-S24). This enzyme catalyzes O-methylation of secondary metabolites and isoprenoid intermediates. Its consistent presence across all *P. pasteri* 2 and absence in *P. pasteri* 1 makes it a candidate marker of the plaque-associated ecotype, though the specific functional contribution to plaque colonization remains to be determined experimentally. Together, these results indicate that *P. pasteri* comprises two ecologically distinct ecotypes–one mucosal-associated, and one plaque-associated that have diverged without major gene-content differentiation, consistent with models of microbial ecotypes driven by regulatory or other non-accessory-gene variation (32). **Distribution of genes in human oral habitats**

Gene-level detection patterns across *Porphyromonas* taxa reinforced the ecological specificity observed at the genome scale. For *P. pasteri* JCM 30531 and *P*. *catoniae* ATCC 51270, gene detection closely mirrored genome-level distributions (Fig. 5A). *P. pasteri* genes were broadly detected on the tongue dorsum with reduced detection in dental plaque, while *P*. *catoniae* genes were predominantly detected in supragingival plaque. In addition to ribosomal and transfer RNAs, which recruited reads across all sites as expected for highly conserved genes, clusters of accessory genes in *P. pasteri* showed cross-site detection, consistent with low divergence among ecotype 1 and 2 subpopulations at the nucleotide level.

**Fig. 5.**
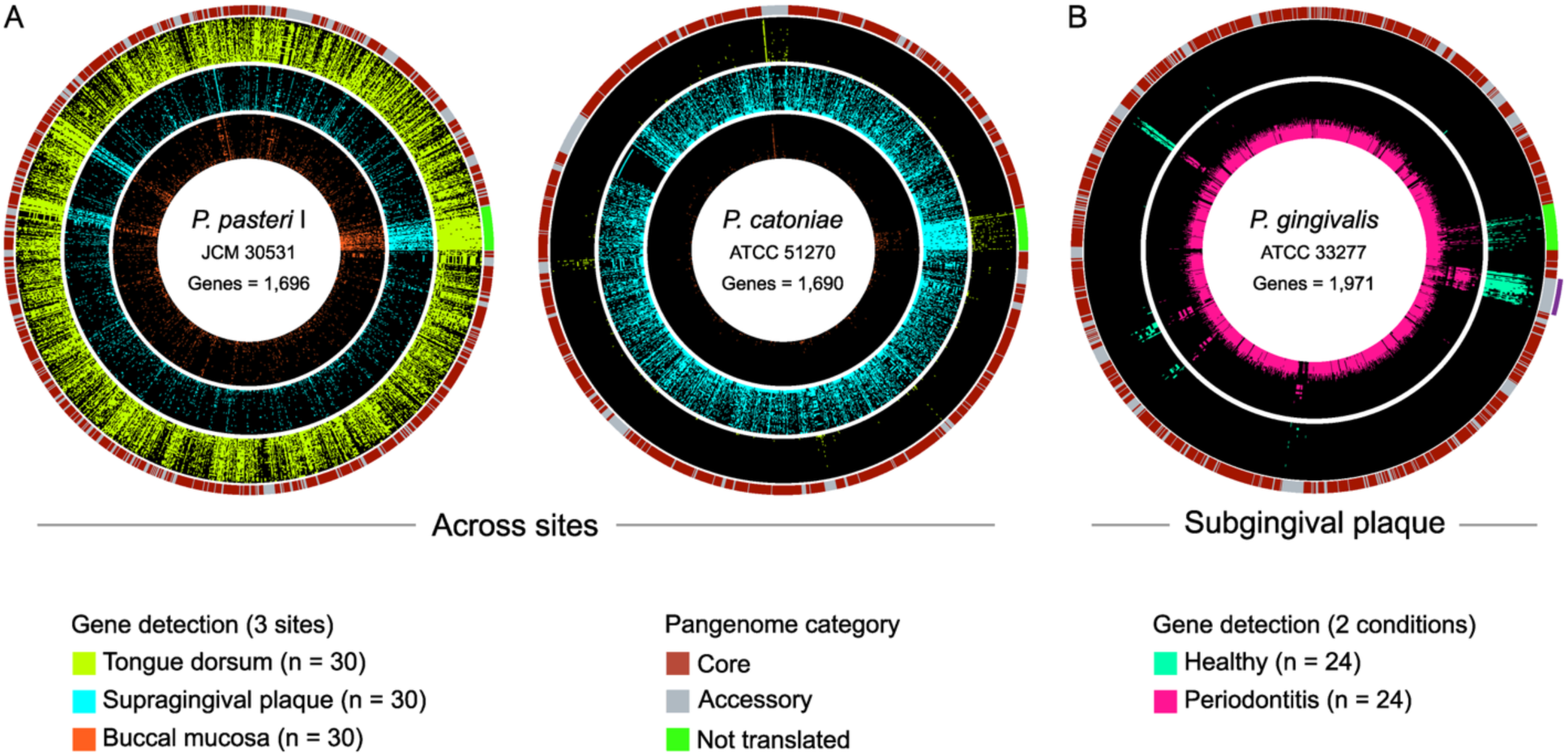
Gene-level detection patterns in health and disease for representative *Porphyromonas* strains. Circular binary plots show detection of predicted genes (gene breadth ≥ 90%) across (A) healthy tongue dorsum (TD; yellow), supragingival plaque (SUPP; cyan), and buccal mucosa (BM; orange) for: *P. pasteri* JCM 30531 and *P*. *catoniae* ATCC 51270. (B) healthy subgingival plaque (SUBP; teal) and periodontitis-associated subgingival plaque (PERIO; pink) for *P*. *gingivalis* ATCC 33277. Black indicates a gene was not detected (gene breadth < 90%). For each genome, the top 30 metagenomes with the highest median gene depth of coverage per site were selected (24 metagenomes for SUBP and 24 for PERIO). Genes were ordered by synteny except for tRNAs and rRNAs and classified as core, accessory, or non-translated (rRNAs, tRNAs), shown in the outermost ring, based on a species-level pangenome. Within each site, metagenomes are rank-ordered by increasing genome breadth of coverage. A purple arc in *P*. *gingivalis* identifies a 44 kb conjugative element.

For *P*. *gingivalis*, genes were broadly detected (gene breadth ≥ 90%) across the subset of periodontitis samples but largely absent from healthy subgingival samples (Fig. 5B), even at a relaxed threshold (gene breadth of coverage ≥ 10%). Examination of virulence-associated loci in *P*. *gingivalis* revealed that gingipain R was consistently detected in periodontitis samples, coinciding with *P*. *gingivalis* genome-level detection, while gingipain K appeared only sporadically, indicating population-level variation in virulence gene content.

A notable exception to the absence in healthy samples of *P. gingivalis* genes was the presence of a discrete 44 kb contiguous block. Inspection of nucleotide-level coverage profiles across this region revealed high depth of coverage spanning the full 44 kb block (Fig. S7), confirming that the locus was present even where the broader *P*. *gingivalis* genome was not detected.

This locus, first found in the *P*. *gingivalis* ATCC 33277 type strain and dubbed CTnPg1 (27, 28), encodes a cohesive set of functions characteristic of a conjugative genomic island, including a Type IV secretion system (VirB4/VirD4), a recombinase, and partitioning and replication proteins, together indicating capacity for conjugative transfer, integration, and stable maintenance. A complete functional annotation of genes within this genomic island and its flanking regions is provided in Table S25. Together, these features define this 44 kb locus as a discrete, conjugative element capable of dissemination across taxa.

Because reads were mapped competitively to all known *Porphyromonas* genomic groups detectable in the human oral cavity (Fig. 3), detection of this locus in healthy samples where *P*. *gingivalis* itself is absent indicates that these genes reside in other members of the oral community. To investigate the evolutionary origin of this region, we examined nucleotide-level similarity of genes within the conjugative element to all genomes in the HOMD. Genes within the genomic island showed high nucleotide identity to homologs distributed across multiple non-*Porphyromonas* oral taxa, predominantly within the order *Bacteroidales* (Table S26). For most island genes, the top matches fell within this family despite divergence at the genus and species level. Similarly, amino acid-level comparisons resulted in concordant patterns of homology (Table S27), supporting a shared evolutionary origin. These island genes occur as accessory gene clusters in a subset of *P*. *gingivalis* strain genomes (Fig. S7) rather than universally across all strains of the species, suggesting that this element is not necessary for pathogenesis. Together, our results indicate that this conjugative element circulates among co-resident taxa in the oral community, independently of the *P*. *gingivalis* genome.

## Discussion

Microbial communities in the human oral cavity are structured by spatially defined ecological niches. An increasing body of research supports the site-specialist hypothesis: that microbes from closely related taxa have adapted to distinct oral sites, and that these patterns can be revealed when communities are analyzed with sufficient resolution (9–13). Our study reinforces this concept, demonstrating that members of the genus *Porphyromonas* exhibit clear site-specialization at both the species and strain levels. By combining metagenomic mapping with pangenomics and gene-level analyses, we connect ecological patterns to functional and evolutionary signatures, thus establishing an ecological framework for understanding evolutionary diversity and microbial niche adaptation.

Establishing a robust genomic reference was essential for interpreting metagenomic ecological data. By integrating genetic content from the pangenome, average nucleotide identity, and phylogenomic analysis, we reconciled species names, corrected misclassifications, and dereplicated near-identical genomes, enabling us to develop a curated set of reference genomes. This curation limits dilution of mapping to redundant genomes, aligns ecological signal to well-defined genomic units, and permits determination of the relative abundance of naturally occurring populations.

Closely related genomic groups often displayed mutual exclusion patterns. Such patterns can arise from competition for the niche. In this concept, similar taxa have the potential to occupy a niche but cannot do so if a member of the group is already present. As a result, each genomic group is found in a subset of individuals. In dental plaque, for example, *P*. *catoniae* and its sister clades alternated dominance across individuals. In diseased subgingiva, *P*. *gingivalis* and *P*. *endodontalis* tended not to be abundant in the same samples. Alternatively, this pattern could be explained by niche partitioning (8), where the two taxa occupy distinct niches, with the niches being of uneven distribution from sample to sample.

At the species level, niche partitioning in *Porphyromonas* is accompanied by divergence in metabolic capacity, consistent with the site specialization of other oral genera. The restriction of *P*. *catoniae* and sister clades to dental plaque correlates with their reliance on exogenous essential nutrients–lacking biosynthetic pathways for cobalamin, biotin, and several amino acids–implying a metabolic interdependence with neighboring taxa in the community, a pattern also observed for *Veillonella*, *Gemella*, and other oral specialists (10, 11). These dependencies fit the Black Queen hypothesis (33), in which community members reduce metabolic burden, relying on co-occurring taxa through gene loss. Conversely, *P*. *gingivalis* possesses the broadest metabolic repertoire in the genus, consistent with the capacity to persist under fluctuating nutrient and host immune conditions without relying on stable community partners. That *P*. *gingivalis* was absent from the majority of periodontitis samples reinforces the polymicrobial synergy model of periodontal disease (31), in which community-level ecological shifts, rather than the presence of any single pathogenic bacteria, drive pathogenesis.

The discovery that *P. pasteri* comprises two ecologically distinct ecotypes without discernible gene content or metabolic differentiation represents a case of habitat divergence. Both phylogenetically defined clades share near-identical functional profiles yet occupy contrasting habitats–one tongue dorsum and mucosal, one dental plaque–consistent with the emergence of bacterial ecotypes through mechanisms beyond accessory gene gain or loss (32). Our competitive mapping results quantitatively extend and confirm the phylogenetic observation by Galtier et al. (24) that *P. pasteri* from dental plaque and tongue dorsum are evolutionarily distinct lineages; while phylogeny can define the existence of subpopulations, competitive mapping can go further by revealing how completely each ecotype permeates different oral environments across individuals–a resolution not achievable from phylogenies alone. The sole enriched function in *P. pasteri* 2–Leucine carboxyl methyltransferase–is a candidate molecular determinant of the plaque ecotype. However, its specific functional role in plaque colonization remains to be determined. More broadly, ecological divergence in the absence of major differences in gene content points to regulatory variation, amino acid variation in adhesin or surface-protein binding sites, or subtle differences in gene expression as the underlying determinants of habitat preference–mechanisms that have been implicated in ecotype divergence in marine bacteria (26).

Although the 44 kb conjugative element is well known in *P*. *gingivalis* ATCC 33277, its detection across oral microbial communities independently of the *P*. *gingivalis* chromosome was an unexpected finding of this study, indicative of horizontal gene transfer. The element encodes the full machinery for conjugative transfer, integration, and maintenance–including a Type IV secretion system, recombinase, and partitioning proteins–alongside defense functions and an efflux transporter that may confer a fitness advantage in healthy subgingival plaque communities. The element’s presence in only a subset of *P*. *gingivalis* strains suggests a dynamic process of horizontal gain and loss. High amino acid identity of gene matches in other species, notably *Prevotella*, indicates that horizontal transfer is not limited to *Porphyromonas* species but can occur across genera within the order *Bacteroidales*, and the element’s presence in healthy samples likely reflects the composition of co-resident taxa, enabling *Prevotella* and related taxa to serve as both donors and recipients.

This pattern of cross-genus element circulation extends a key concept in microbial ecology: that the accessory genome of a community can function as a distributed genetic source whose ecological fate is partially decoupled from any single host lineage (34, 35). Here, we extend this concept to conjugative elements capable of carrying adaptive traits across co-resident taxa at greater evolutionary distances than traditional boundaries would suggest, and doing so in a habitat-specific manner that is detectable at scale through competitive metagenomic read recruitment.

In conclusion, our study provides a genomic and ecological framework for understanding *Porphyromonas* site-specialization in the human oral microbiome. By integrating genomic content, ecological distribution, and functional potential, we characterized how evolutionary divergence, metabolic interdependence, and mobile genomic elements shape microbial population structure in health and disease.

## Methods

All analyses were performed using the Anvi’o (v8) platform (36, 37) with Python (v3.10.15) and R (v4.4.1) under the Anaconda platform (conda v24.11.3).

### Porphyromonas genomic set

All publicly available *Porphyromonas* genomes were downloaded from the National Center for Biotechnology Information (NCBI) RefSeq database using the ‘dataset’ command-line tool (v17.2.0) with the taxon flag ‘*Porphyromonas*’ on June 23, 2025 (38). To maximize representation of the genus, we kept genomes annotated at both species and genus levels (e.g., *Porphyromonas* sp.), as well as those originating from metagenome-assembled genomes (MAGs), yielding 377 genomic assemblies (Table S1).

### Quality control

Completeness and contamination were estimated using lineage-specific marker genes with CheckM (v1.2.3) (39) and pre-calculated machine-learning-based markers with CheckM2 (v1.1.0) (40). Further, we evaluated completion and redundancy of universal bacterial marker genes (Anvi’o Bacteria_71) (41). Genomes were retained if ≥90% complete and <5% contaminated for both CheckM and CheckM2, and for universal marker genes ≥70% completion with <10% redundancy was required (Tables S2-S4). After filtering, 343 high-quality genomes remained.

### Human oral genomic set

A genus-level phylogeny was constructed from 59 Bacteria_71 marker genes extracted, individually aligned with MUSCLE (v3.8.1551) (42), concatenated, and trimmed with trimAl (v1.4) to remove positions with ≥ 50% gaps (Table S5) (43). A maximum-likelihood phylogeny was inferred with IQ-TREE (v2.4.0) (44) with the Whelan and Goldman (WAG) amino acid substitution model and 1,000 bootstrap replicates (UFBoot) (45) rooted with *Bacteroides pyogenes* NCTC11853, *Prevotella melaninogenica* “ATCC 25845”, *Tannerella forsythia* “ATCC 43037”, and *Tannerella serpentiformis* W11667, and visualized in R using the ggtree package (46). Taxonomic assignment was performed with the Genome Taxonomy Database Toolkit GTDB-Tk (v2.4.1) (47, 48) using release r226 (Table S6). Human oral clades were identified by cross-referencing GTDB assignments with the Human Oral Microbiome Database (HOMD; v4.0) and NCBI ecological metadata (isolation source and host). Species-ambiguous habitat assignments were resolved by mapping reads from oral and non-oral human body sites to type strain genomes (Table S8). We therefore retained clades with named human oral species and clades with unnamed genomes (*P*. sp.) supported with human oral metadata.

### Genome contigs database and annotation

Genomes were processed with ‘anvi-script-reformat-fasta’ and ‘anvi-gen-contigs-database’ to keep contigs ≥ 200 bp, replace non-canonical bases with the letter ‘N’, and predict open-reading frames (ORFs; hereafter genes) with Prodigal (v2.6.3) (49). Genes were annotated with NCBI Cluster of Orthologous Genes (COG 20) (50) via Diamond (v2.1.11) (51), Pfam (v37.2) (52) KEGG KOfams and modules (v2023-09-22 snapshot) (53, 54), and CAZymes (v13) (55) via HMMR (v3.4). Ribosomal genes, Bacteria 71 marker genes (41), and tRNAs (tRNAscan-SE, v2.0.7) (56) were identified with ‘anvi-run-hmms’ and ‘anvi-scan-trnas’.

### Pangenome construction

Pangenomes were constructed in Anvi’o following previously described steps (25). We used BLASTp (v2.16.0+) for all pairwise amino-acid sequence comparisons, grouped by Markov Clustering Algorithm (MCL; v22-282; inflation parameter = 10 and minbit = 0.5) (57). The resulting homologous gene clusters were used to group genomes and gene clusters hierarchically using Euclidean distance and Ward linkage to visualize patterns of gene-content variation across genomes.

### Average Nucleotide Identity and Tanglegrams

Pairwise ANI was calculated using ‘anvi-compute-genome-similarity’, which implements pyANI (v2.11) (58) with the ANIb method (BLASTn; v2.16.0+) and a minimum alignment fraction of ‘0’. To compare ANI-based and phylogenomic clustering, we identified single-copy core genes from the pangenome, concatenated their sequences, aligned and trimmed gap positions (>50% gaps), and inferred a maximum-likelihood inference with IQ-TREE as described above. Tanglegrams were constructed by co-visualizing the result tree with the ANI-based dendrogram, minimizing line entanglement to highlight concordant genome placements using ‘dendextend’.

### Reference genomes for metapangenomics

The 191 human oral genomes were dereplicated at 98% ANI using ‘anvi-dereplicate-genomes’ with a simple greedy algorithm, minimum alignment fraction of 25%, and centrality for pre-selected reference genomes. For each dereplication cluster, we manually select by preference of isolates over MAGs, type strains status, and overall genome quality, yielding 84 reference genomes (Table S9).

### Metagenomic sample set

Healthy human oral metagenomes from nine sites were obtained from the Human Microbiome Project (HMP) portal (3). Periodontitis metagenomes (n = 24) were kindly provided by Scott T. Kelly (59). HMP samples were downloaded via SRS identities from NCBI (1,275 metagenomes from 220 individuals). Gingival samples were further reclassified as keratinized and attached gingiva, supragingival plaque, and subgingival plaque by manual metadata review. Metagenomes were quality-filtered following recommendations by Minoche, Dohm, and Himmelbauer (60) with the program ‘iu-quality-minoche’ (https://github.com/merenlab/illumina-utils) (61) and kept those with ≥1 million paired-end reads. Human DNA was removed by the HMP team according to their guidelines. The final dataset comprised 1,218 healthy metagenomes (25,264,824,141 paired-end reads; nine sites; 220 subjects) and 24 periodontitis-associated metagenomes (295,490,295 paired-end reads; 24 subjects).

### Competitive metagenomic mapping

Reads were mapped to the 84 dereplicated reference genomes using Bowtie2 (v2.5.4) with the ‘--end-to-end’, ‘--very-sensitive’, and ‘--no-unal’ flags (62). SAM files were binarized, sorted, and indexed using samtools (v1.21) (63). Coverage metrics were calculated with ‘anvi-profile’ and merged by site (anvi-merge). Breadth of coverage and depth of coverage metrics were summarized in tables with ‘anvi-summarize’ with the ‘--calculate-q2q3-carefully’ flag (Anvi’o v8-dev). A genome was considered detected if the breadth of coverage was ≥ 50% (at least half the nucleotides covered by ≥ 1 read). Relative abundance was calculated as the Q2-Q3 mean depth of coverage for each genome divided by the sum of Q2-Q3 mean depths of coverage across all genomes in a sample; species-level relative abundance was the sum across all genomes in a genomic group.

### Pairwise mutual exclusion analysis

To quantify pairwise mutual exclusion among *Porphyromonas* taxa, we evaluated complementary presence-absence and abundance-based metrics. Presence-absence associations were assessed using Fisher’s exact test (odds ratios). Abundance relationships among co-detected samples were evaluated using the centered log-ratio (CLR) transformation and Spearman correlation. Dominance direction was determined by the Wilcoxon signed-rank test on CLR differences between co-occurring pairs. P-values were adjusted for multiple testing using Benjamini-Hochberg false discovery rate (FDR).

### Metabolic profiles and functional enrichment analysis

We used ‘anvi-estimate-metabolism’ to assess the completeness of metabolic pathways across each dereplicated genome. Using a ‘pathwise’ approach, in which all possible path combinations in a module are explored, we kept the one with the highest completeness score. For a pathway to be ‘complete’, at least 75% of the necessary enzymes should be present. Functional and metabolic enrichment analyses were estimated by linking genomes to a site (or condition in the case of periodontitis), and then we quantified the distribution of functions or modules in each group of genomes. Enrichment test was assessed with a Generalized Linear Model with a logit linkage function to compute an enrichment score and p-value for each function or module–using the ‘q-value’ package in R. This evaluates, for every function or module, whether the occurrence in the genomes is greater than the expected occurrence under uniform distribution. We considered a function or module to be truly enriched if the adjusted q-value was less than 0.05 and present in at least 50% of the group members.

### Validation of *P. pasteri* ecotypes

Non-RefSeq *P. pasteri* genomes and assemblies lacking species designation were retrieved from GenBank (n = 491). From this set, seven high-quality genomes passing all quality thresholds, GTDB taxonomic annotation, and phylogenomic analysis were retained (Table S18 and S19) and combined with the 18 high-quality RefSeq *P. pasteri* genomes. Dereplication at 98% ANI resulted in 24 representative genomes. Competitive mapping was performed using the 50 metagenomes per site with the highest *Porphyromonas*-mapping read counts across tongue dorsum (TD, n = 50), supragingival plaque (SUPP, n = 50), and buccal mucosa (BM, n = 50). Relative abundance was estimated as before, using the careful Q2-Q3 mean depth of coverage.

### Gene-level distribution analysis

To produce gene-level plots with syntenic organization, we ran ‘anvi-interactive’ in ‘gene-mode’, which computes a profile database with gene-level metrics. For each genome, we focused on the top 30 metagenomes with the highest median gene depth of coverage per site. We defined a gene as ‘detected’ if at least 90% of its nucleotides were covered 1X. Gene categories (core, accessory, non-coding) were identified from species-level pangenomes after removal of duplicates (same strain deposited in with different ID–e.g., ATCC and NCTC). We used ‘anvi-get-split-coverage’ to extract per-nucleotide coverage for the region of interest.

### HOMD BLAST analysis

To infer taxonomic origin of conjugative element genes, we performed nucleotide- and amino acid-level similarity searches against the HOMD. Searches were conducted using the HOMD Genomic BLAST Server (https://blast.homd.org/genome_blast/) with an e-value of 1×10^−5^ and a maximum of 20 target sequences using the NCBI All Genomic DNA or All Proteins Annotated from the HOMD Genomes database v11.02. For each query, we kept only the top three hits for downstream analysis.

## Acknowledgments

We are grateful to A. Murat Eren and the Anvi’o team for their assistance, consultation, and support.

## Funding

NIH NIDCR DE030136 (GGB, JMW, and FED), NIH NIDCR DE016937 (JMW and FED), and Forsyth Collaborative Pilot Grant FSI_CP04 (GGB and FED).

## Author contributions

Conceptualization: JTM, JMW, GGB Methodology: JTM

Investigation: JTM Data Curation: JTM Visualization: JTM

Funding Acquisition: FED, JMW, GGB Project Administration: JMW, GGB Supervision: JMW, GGB

Writing – original draft: JTM

Writing – review & editing: JTM, FED, JMW, GGB, KMK

## Competing interests

The authors declare no conflicts of interest.

## Data Availability

All data in this study are publicly available.

Genomes were obtained from NCBI: https://www.ncbi.nlm.nih.gov/datasets/genome/

Metagenomes were obtained from the Human Microbiome Project portal: https://portal.hmpdacc.org/

Code availability: https://github.com/juliantom/Metapangenomics_Porphyromonas_2025_06_23

## Supplementary Materials

Extended Data (text and Figs. E1 to E3)

Figs. S1 to S7

Tables S1 to S27

## Extended Data

### Phylogeny of *Porphyromonas* taxa

To establish a curated framework for identifying human oral genomes within the genus *Porphyromonas*, we first examined the evolutionary relationships among all 343 quality-filtered publicly available genomes. Many were deposited as *P*. sp., and numerous metagenome-assembled genomes (MAGs) lacked standardized species designations, creating ambiguity that complicated comparative analyses. We resolved these inconsistencies by constructing a phylogeny of the *Porphyromonas* genus from universal bacterial marker genes (Extended Data Fig. E1). To standardize taxonomic assignments, we used the Genome Taxonomy Database Toolkit (GTDB-Tk; release r226) to provide species-level assignments independent of historical nomenclature. The resulting tree revealed well-supported monophyletic clades representing coherent evolutionary groups comprising genomes with recognized species names, Human Microbial Taxon (HMT) designated taxa, or unnamed *P*. sp. genomes. This phylogenetic framework thus provided a foundation for taxonomic harmonization and the identification of candidate human oral genomes.

We next annotated each genome according to its isolation metadata from NCBI and assigned genomes to one of three putative categories: human oral, human non-oral, or animal–as shown in the ‘Ecology layer’ of Extended Data Fig. E1. Because isolation metadata may reflect sampling bias rather than true ecological habitat, taxa were considered potentially human oral when at least one genomic member had a documented origin from the oral cavity or upper respiratory tract. Using this criterion, human oral taxa comprised: *P*. *gingivalis*, *P*. *endodontalis*, *P*. *catoniae*, *P*. sp. HMT-275, *P*. sp. HMT-278, *P*. *pasteri*, and several unnamed *P*. sp. clades–labeled according to the closest named species as indicated in the ‘Taxa’ layer of Extended Fig. E1. Non-oral human *Porphyromonas* were included in the phylogeny to provide ecological context. Notably, human oral taxa are phylogenetically closer to animal taxa than to *Porphyromonas* isolated from other human body sites. *Porphyromonas asaccharolytica* (HMT-547) and *P*. *uenonis* (HMT-785) are listed in HOMD (v4.1) with ‘uncertain’ or ‘primarily vaginal’ habitats, and along with *P*. *vaginalis* they group in the same branch. The genome *P*. sp. ‘31_2’ (GCA_000712235.1) deposited as *Porphyromonas* was classified as *Parabacteroides distasonis* and showed relatedness to *Tannerella*. This isolation metadata-based approach generated a set of candidate human oral genomes pending ecological validation.

### Ecological validation of named taxa across the human body

To assess whether candidate human oral taxa recruited reads from oral habitats, we performed competitive mapping of shotgun metagenomic sequence data to quality-filtered type strain genomes. We used metagenomic data from the Human Microbiome Project which sampled four human body sites in healthy individuals: the mouth, stool, vagina, and skin. Within these sites, data were available from nine oral subsites, four skin subsites, and three vaginal subsites.

We also included data from a study of subgingival plaque in periodontitis (Extended Data Fig. E2). This analysis confirmed that taxa identified as oral candidates based on phylogeny and metadata consistently recruited reads from oral metagenomes.

*Porphyromonas uenonis* was detected primarily in vaginal samples, but also at low prevalence in oral sites. *Porphyromonas asaccharolytica* and *P*. *vaginalis* also predominantly recruited reads from vaginal metagenomes, but had negligible recruitment from oral metagenomes*. Porphyromonas bennonis*, *P*. *somerae*, and *P*. *miyakawae* were not detected across any surveyed body site. Notably, despite its historical role as the reference species of the genus, *P*. *asaccharolytica* showed no oral detection, indicating it is unlikely to be a constituent of the oral microbiome. The detection of *P*. *uenonis* in both vaginal and some oral samples suggests that some *Porphyromonas* taxa can occupy multiple mucosal niches rather than being restricted to a single body site.

### A phylogenetically and ecologically defined oral set

By constructing phylogenomic relationships, resolving taxonomic identity and validating the ecological distribution of named species across body sites, we delineated a human oral subset of genomes for the genus *Porphyromonas*. This set includes both named taxa with established type strains and as yet unnamed clades representing potential novel taxa. This set of 191 genomes forms the curated reference set of genomes for the metapangenomic analyses in the main text.

### Pangenome of 191 human oral *Porphyromonas* genomes

Pangenome analysis of all 191 human oral *Porphyromonas* genomes prior to dereplication (Extended Data Fig. E3) confirmed the existence of coherent phylogenetic groups but also revealed within-group structure that dereplication, by design, compresses.

*Porphyromonas gingivalis* was represented by 99 genomes before dereplication, reflecting high availability of isolates for this disease-associated taxon, but collapsed to a single representative after dereplication, consistent with its extremely low intraspecies genome diversity.

*Porphyromonas pasteri* had only one genome eliminated by dereplication, which corresponded to the *P*. *bobii* strain ‘EMRHCC 6C’ which was deposited twice. Overall, the 12 color-coded genomic groups shown in Extended Data Fig. E3 represent 7 named human oral species (see main text Fig. 1) together with their within-group structure (*P. catoniae*, 3 clades; *P*. sp. HMT-275, 2 clades; *P. endodontalis*, 2 clades; and *P. uenonis*, 2 clades).

Multiple strains, including type strains, have been independently sequenced and deposited by different culture collections; these duplicated assemblies are visible in the pre-dereplicated pangenome. For example, *P*. *endodontalis* ATCC 35406 = NCTC13058 = FDAARGOS 1506 has three independent entries. The inclusion of such duplicates in the pre-dereplicated set enables a quality check on the resulting pangenome. Sequencing of genomes of the same strain multiple times is expected to produce near-identical results, thus serving to measure the accuracy of genomic sequencing. Subtle gene-content differences among duplicates are expected and likely reflect sequencing error rate, assembly workflow, and cultivation passage.

For *Porphyromonas* as a whole, the majority of gene clusters in the pre-dereplicated pangenome carry at least one functional annotation across COG, Pfam, KOfam, or CAZyme databases, indicating that comparative metabolic and functional analyses are well-supported by the available genomic information.

**Extended Fig. E1.**
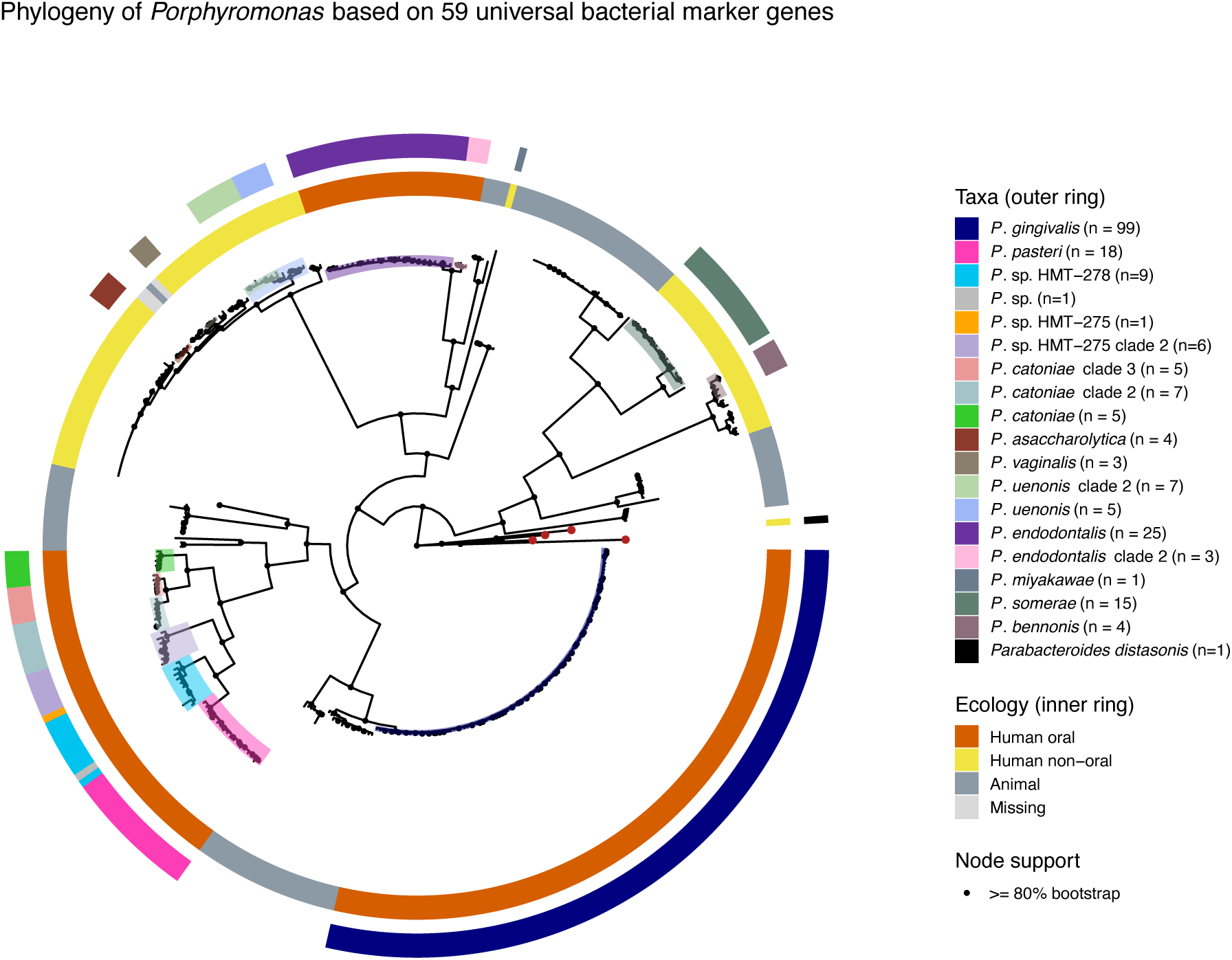
Phylogenomic framework of the genus *Porphyromonas*. Maximum-likelihood phylogeny of 343 quality-filtered *Porphyromonas* genomes inferred from concatenated amino-acid sequences of 59 conserved bacterial marker genes (Bacteria_71 set). Branch lengths represent amino-acid substitutions per site. Node support was assessed using 1,000 bootstrap replicates (UFBoot); only ≥80% are shown (black dots). The tree is rooted using outgroup genomes from *Bacteroides pyogenes*, *Prevotella melaninogenica*, *Tannerella forsythia*, and *Tannerella serpentiformis* (shown with red dots at the tips). Metadata annotations are indicated in the two rings. The outer ring denotes names of curated human species-level taxa along with the number of genomes in each taxon. Names are not shown for animal taxa. The inner ring denotes the ecology (host and body site) for curated genomes, classifying them as human oral, human non-oral, animal, or missing isolation metadata.

**Extended Fig. E2.**
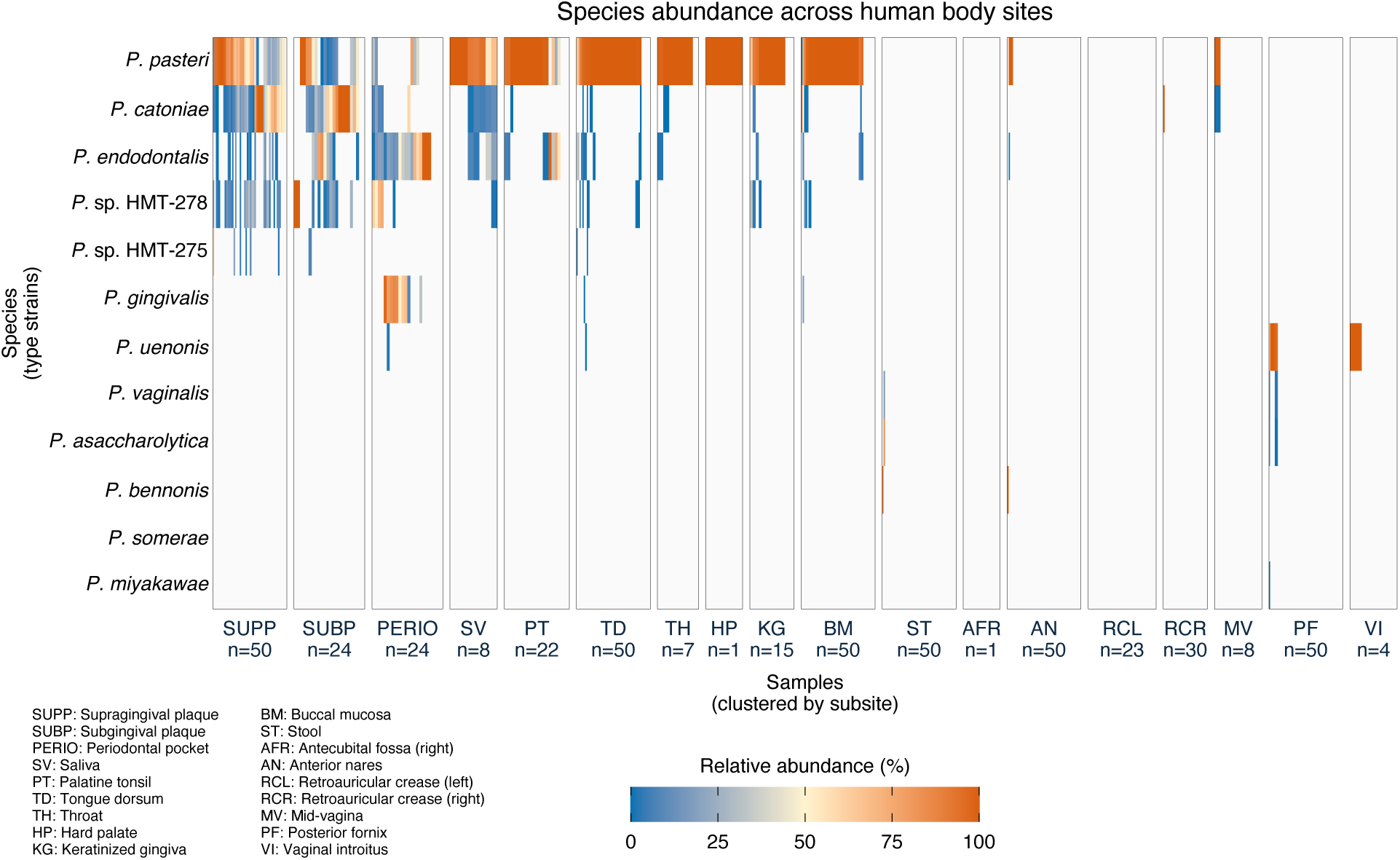
**Distribution of *Porphyromonas* type strains across human body sites**. Relative abundance of type strains representing 12 named *Porphyromonas* species, expressed as a fraction of total *Porphyromonas* abundance, following competitive mapping to up to 50 metagenomes per body subsite from the Human Microbiome Project healthy cohort. Relative abundance is expressed as the fraction of the Q2-Q3 mean depth of coverage for each genome within a given sample. Body subsites are ordered left to right as oral (SUPP, SUBP, PERIO, SV, PT, TD, TH, HP, KG, BM), stool (ST), skin (AFR, AN, RCL, RCR), and vagina (MV, PF, VI). Samples within each subsite are hierarchically clustered by Bray-Curtis dissimilarity with Ward linkage. Light grey indicates no detectable abundance.

**Extended Fig. E3.**
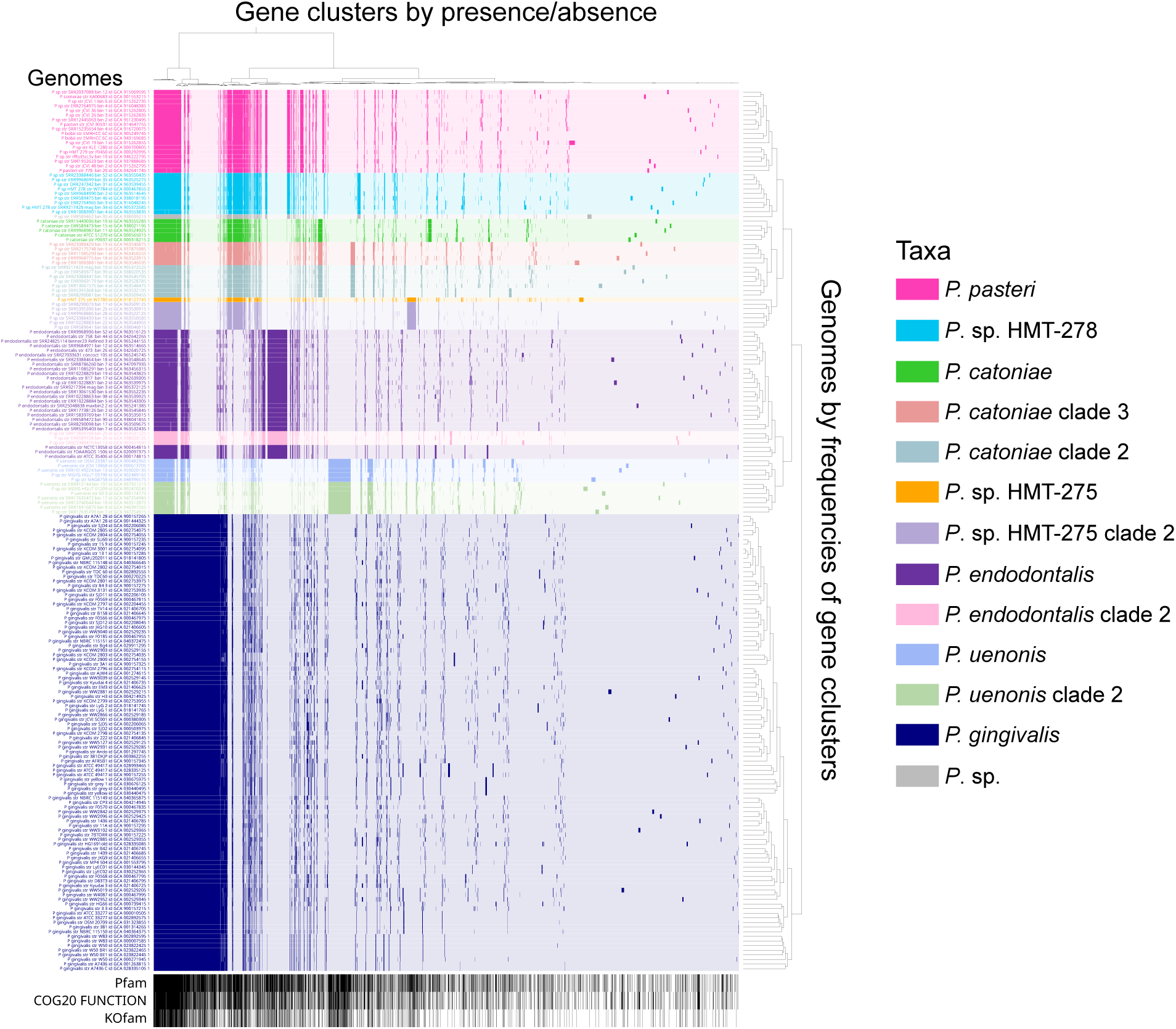
**Pangenome of human oral *Porphyromonas* genomes prior to dereplication (n = 191)**. All 191 high-quality human oral *Porphyromonas* genomes are organized by gene content. Predicted open-reading frames (ORFs) were translated to amino-acid sequences, compared using BLASTp, and grouped into homologous gene clusters (vertical bars) via Markov Clustering Algorithm (MCL; inflation =10). Genomes are hierarchically clustered by gene-cluster frequencies (dendrogram to the right) and are color-coded by genomic group. Black bars at the bottom indicate gene clusters with associated functional annotation.

**Fig. S1.**
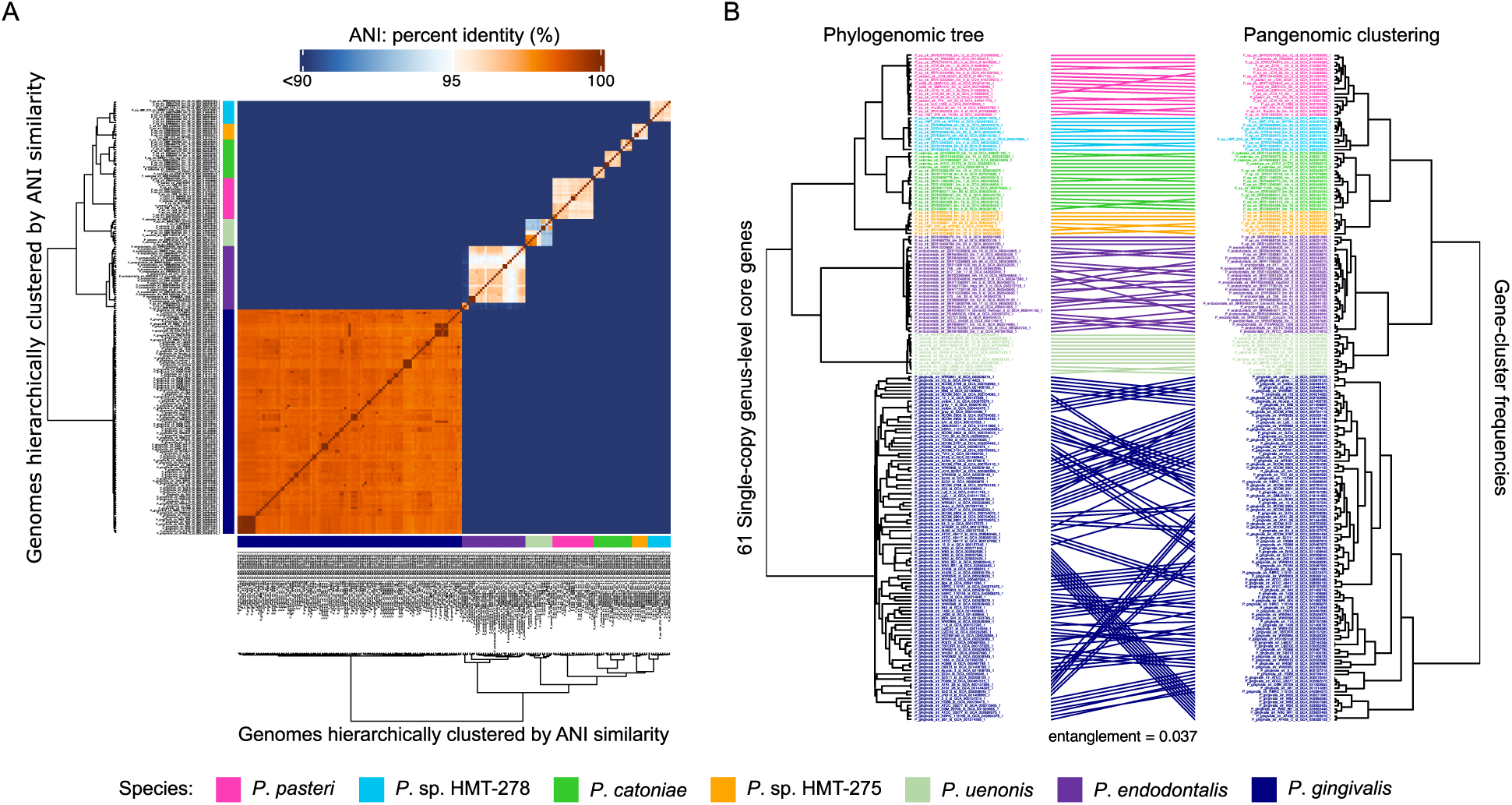
ANI heatmap and tanglegram for human oral *Porphyromonas* genomes. (A) Pairwise ANI heatmap of 191 high-quality human oral *Porphyromonas* genomes reveals that most species show strong intraspecies similarity (>95%) and sharp interspecies boundaries. Values range from 90% (blue) to 100% (red), with a midpoint 95% (white). Genomes are hierarchically clustered by ANI values. (B) Tanglegram comparing a maximum-likelihood phylogenomic tree (left; 61 single-copy core genes, WAG model, 1,000 bootstraps) and a pangenomic tree (right; derived from gene cluster frequency data). Matching genomes are connected by lines arranged to minimize entanglement; labels and lines are color-coded by genomic group designation.

**Fig. S2.**
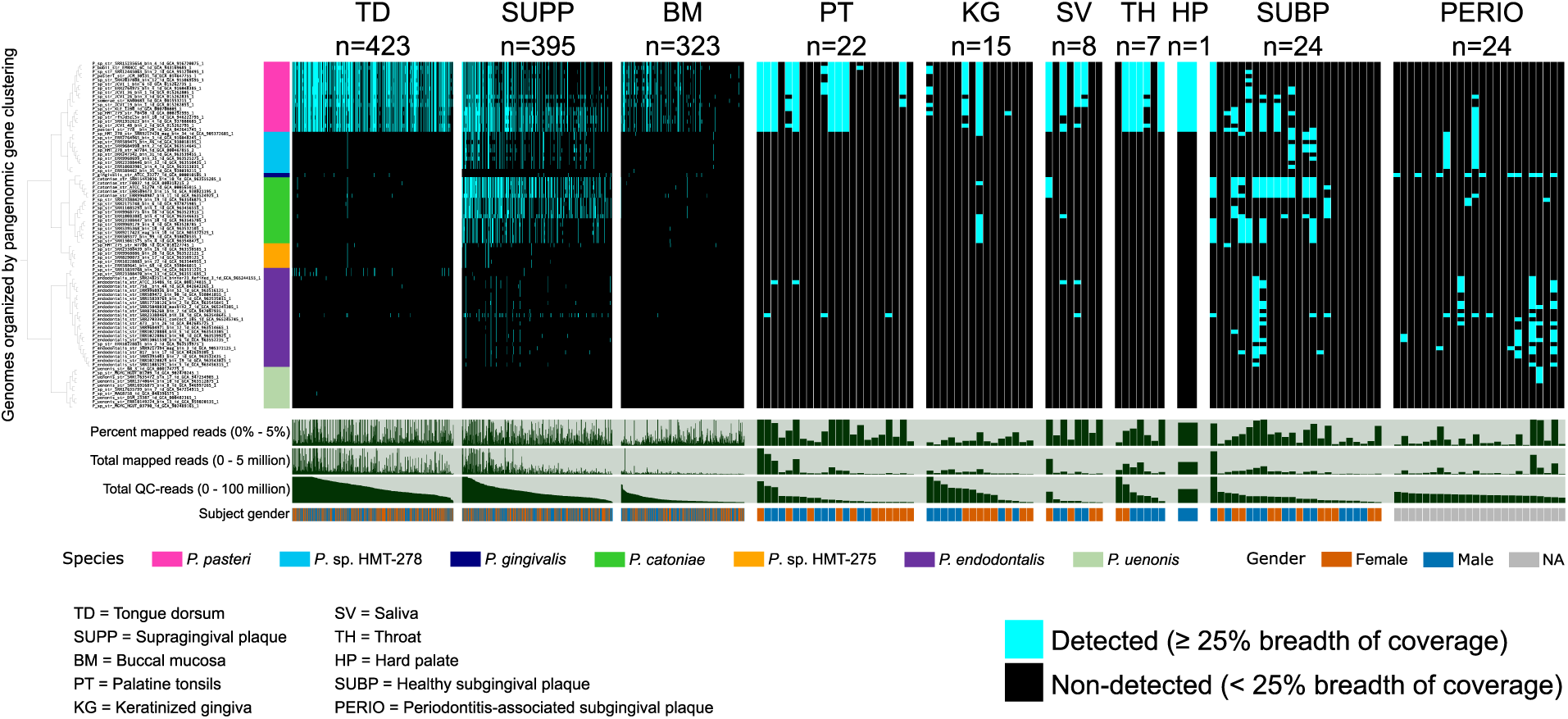
Binary detection of reference *Porphyromonas* genomes across oral sites. Following breadth of coverage, genomes were considered detected (cyan) in a sample if ≥ 25% of nucleotides had at least one mapped read. Detection is shown as a binary presence/absence plot for 84 reference genomes across the 1,242 human metagenomes from nine healthy oral sites and one periodontitis-associated site. Genomes are ordered by gene cluster frequencies (as in Fig. 1); metagenomes are grouped by site and sorted by decreasing read count.

**Fig. S3.**
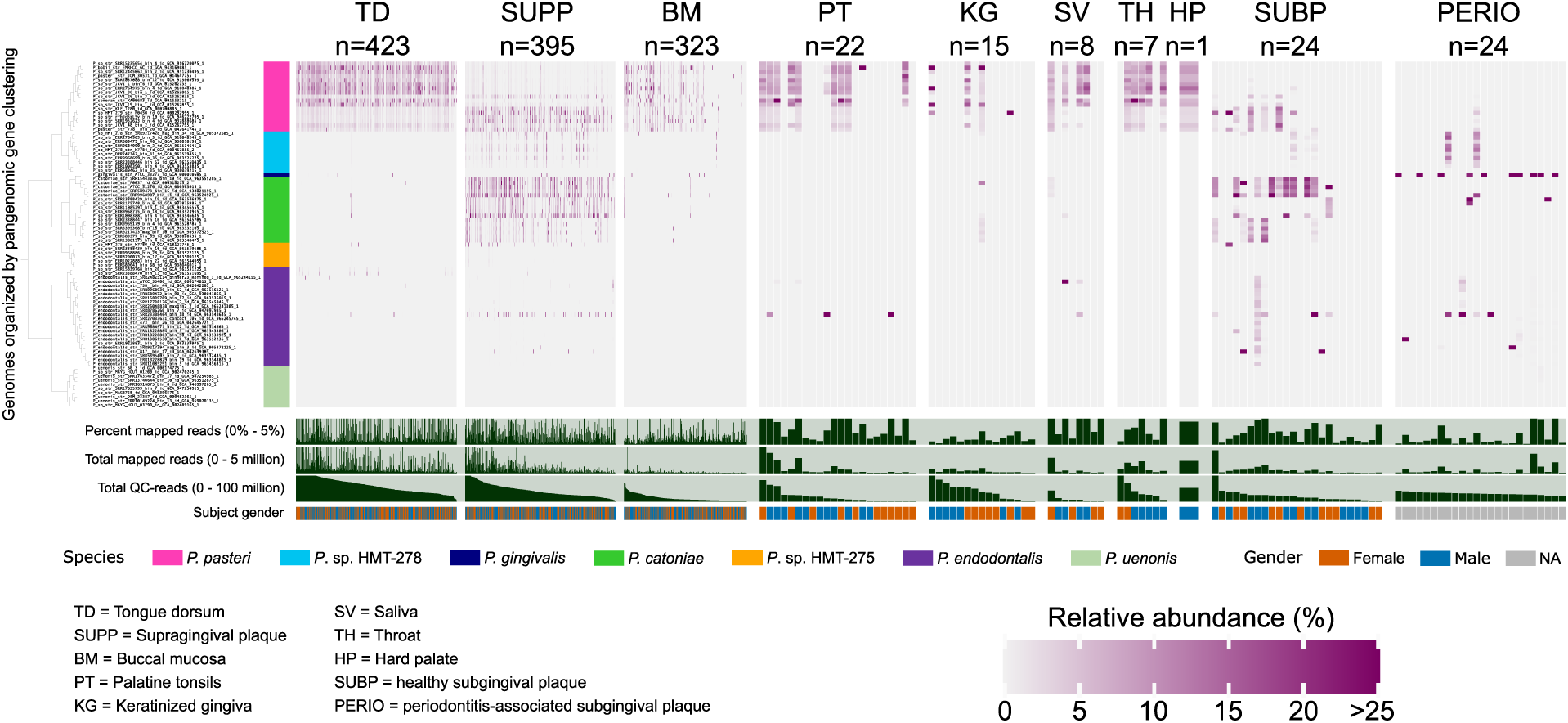
Genome-level relative abundance heatmap. Relative abundance values range from 0% (black) to 25% (magenta) and were calculated as the genomic Q2-Q3 mean depth of coverage divided by the total Q2-Q3 mean depth of coverage across all *Porphyromonas* genomes in a sample. Genomes and metagenomes are ordered as in Fig. S2.

**Fig. S4.**
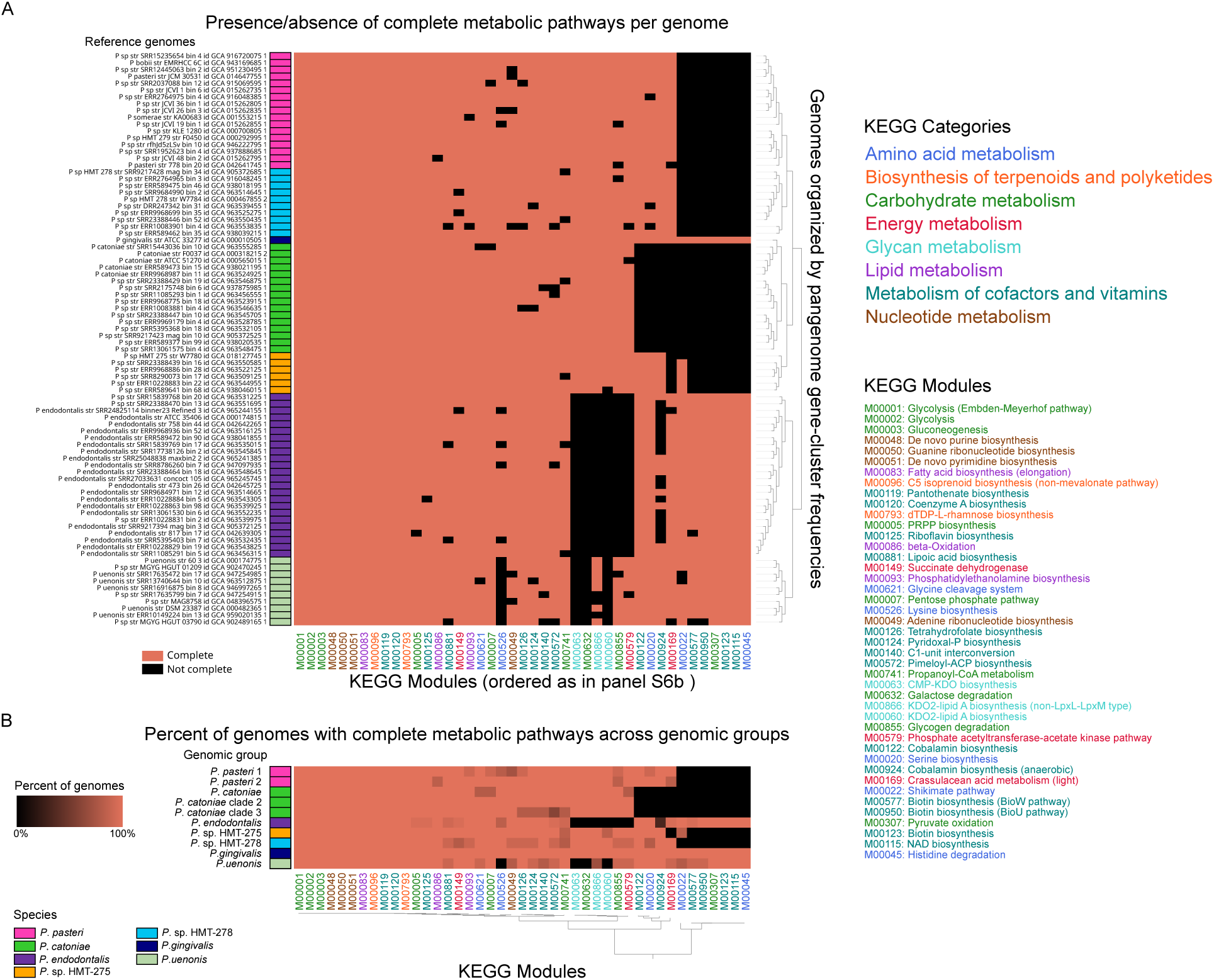
Metabolic completeness across *Porphyromonas* reference genomes and genomic groups. A) Binary plots showing the presence or absence of complete metabolic pathways in 84 reference genomes. A pathway was considered complete if ≥ 75% of the required KEGG module enzymes were present. B) Heatmap showing the percentage of genomes, within genomic groups, in which each metabolic pathway is complete. Modules are hierarchically clustered by Euclidean distance and Ward linkage. Genomic groups have the same order as in Fig. 3.

**Fig. S5.**
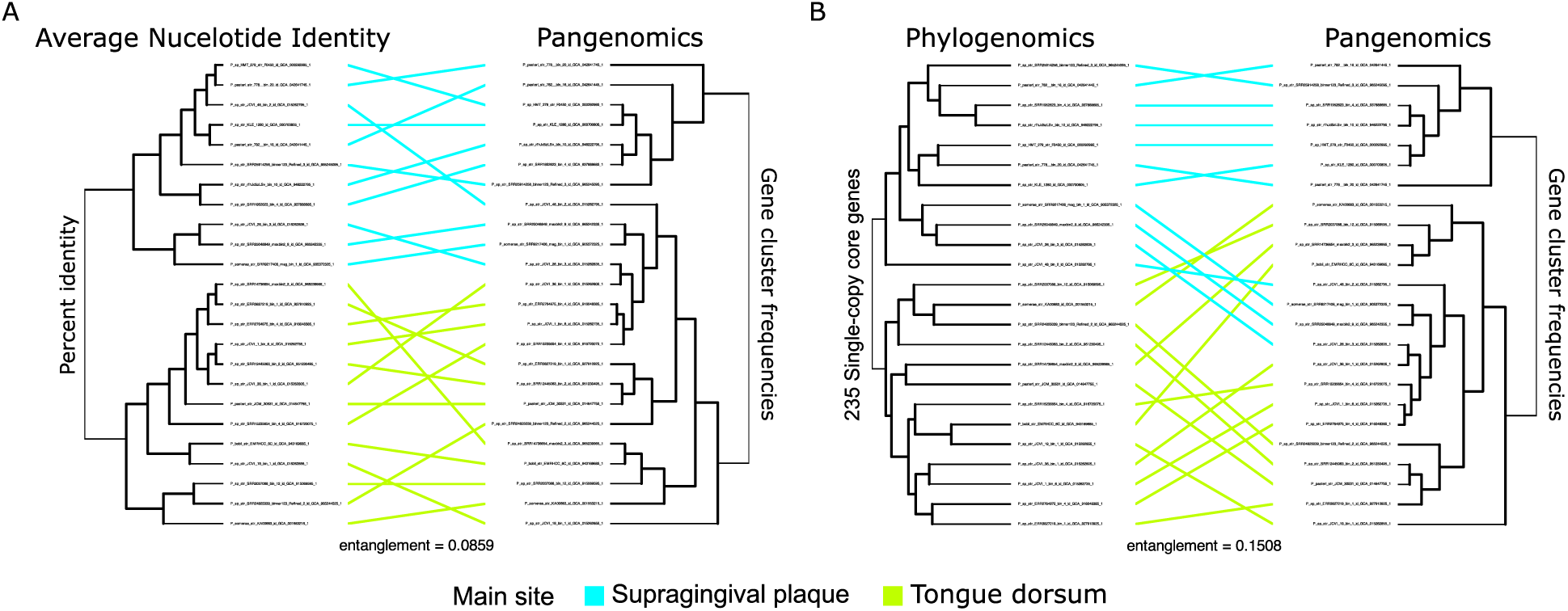
Comparison of *P. pasteri* ecotypes clustering by phylogeny, pangenome, and ANI. Two tanglegrams compare genome clustering based on (A) ANI vs pangenomic analyses, (B) phylogenomic versus pangenomic analyses. Genome labels and connecting lines are color-coded by the primary oral site: tongue dorsum or supragingival plaque. ANI and phylogenomic trees are from analyses presented in Fig. 4. A *P*. *pasteri* pangenome was constructed to generate a gene-cluster frequencies dendrogram.

**Fig. S6.**
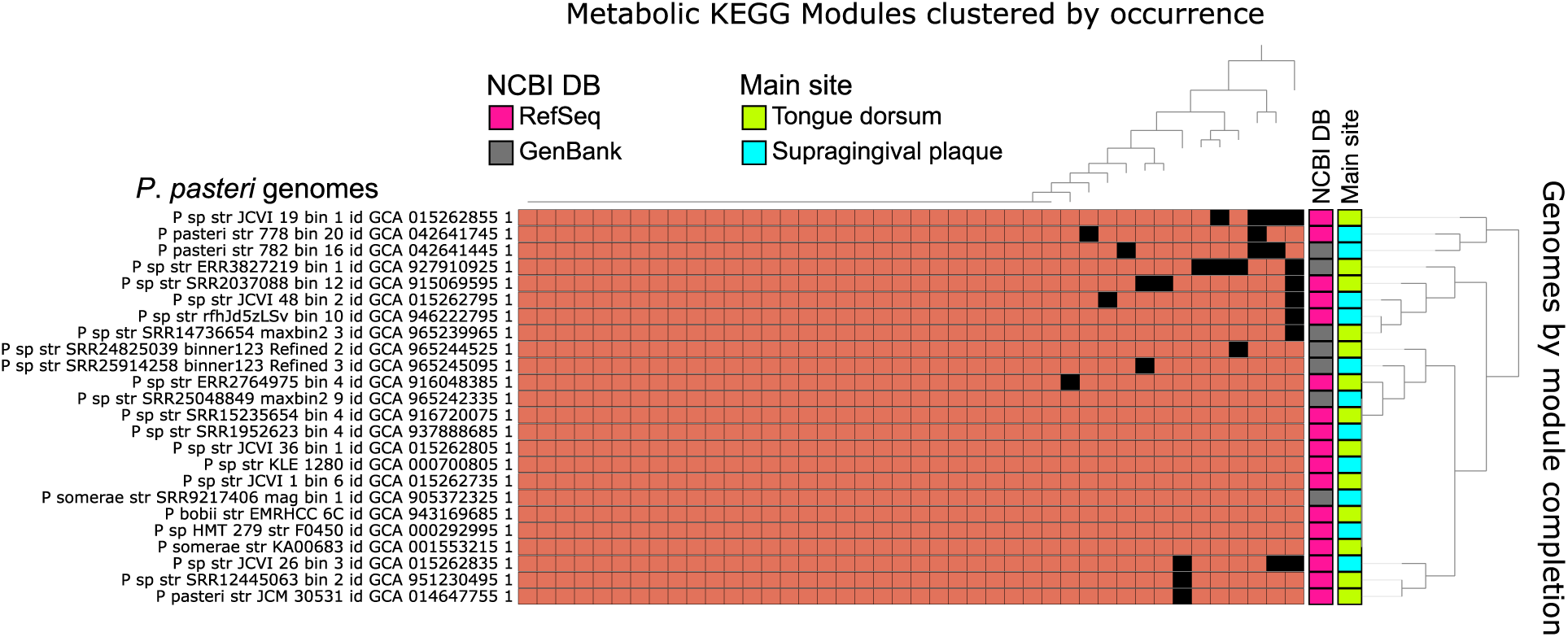
Metabolic capacity of *P. pasteri* ecotypes. Binary plot showing the presence or absence of complete metabolic pathways (≥75% of required KEGG module enzymes). Across *P. pasteri* genomes with distinct site preference: tongue dorsum or supragingival plaque. Labels indicate site preference and NCBI database source (RefSeq or GenBank). Clustering of genomes by module, using Euclidean distances and Ward as linkage, showed limited differences among the specialists.

**Fig. S7.**
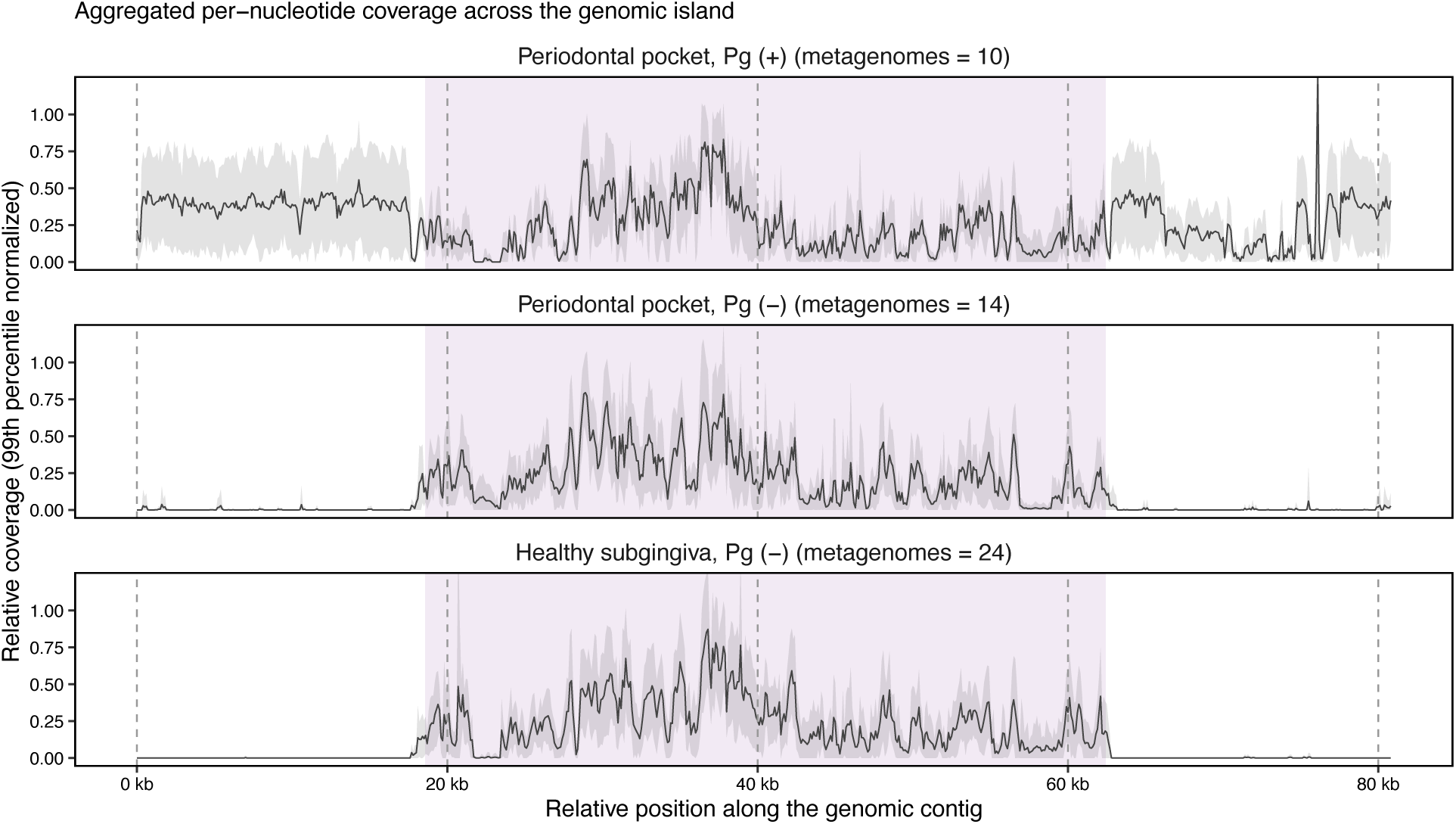
Metagenomic coverage distribution across a 44 kb mobile conjugative element of *P*. *gingivalis*. Per-nucleotide coverage of an 80 kb genomic region containing a mobile conjugative element in *P*. *gingivalis* ATCC 33277 across subgingival plaque metagenomes shows significant coverage despite the absence of the bacterial chromosome. Per-nucleotide coverage for each sample was normalized to the 99^th^ percentile. Normalized coverage was then aggregated across samples within three groups: periodontitis-associated with *P*. *gingivalis* detected (metagenomes = 10), periodontitis-associated with *P*. *gingivalis* not detected (metagenomes = 14), and healthy subgingiva (metagenomes = 24). The group mean is shown as a black line, and the shaded ribbon indicates ± 1 standard deviation across metagenomes. A purple transparent box indicates the relative position of the mobile conjugative element.

**Table S1. Collection of 377 RefSeq *Porphyromonas* genomes.** Two-column table listing the Assembly Accession numbers and corresponding internal genome identifiers for RefSeq *Porphyromonas* genomes available in NCBI (n = 377; date of download: June 23, 2025).

**Table S2. Evaluation of completeness and contamination of the 377 *Porphyromonas* genomes using CheckM.** Columns A-N contain the CheckM (v1.2.3) quality report for each genome. Column O indicates whether each genome passed or failed the quality thresholds of ≥ 90% completeness and < 5% contamination.

**Table S3. Evaluation of completeness and contamination of the 377 *Porphyromonas* genomes using CheckM2.** Columns A-O contain the CheckM2 (v1.1.0) quality report for each genome. Column P indicates whether each genome passed or failed the quality thresholds of ≥ 90% completeness and < 5% contamination.

**Table S4. Assessment of completion and redundancy of marker genes for the 377 *Porphyromonas* genomes.** Columns A-F contain anvi-estimate-genome-completion report of universal bacterial marker genes (Bacteria_71 set). Column G indicates whether genomes passed or failed completion ≥70% and redundancy <10%.

Table S5. Universal bacterial marker genes in the 343 quality-controlled *Porphyromonas* genome collection for phylogenomic analysis. The table presents the 71 genes from the Bacteria_71 universal bacterial marker set, indicating the ones selected for phylogeny analysis. Column A shows the gene names, and Column B indicates whether each gene was identified in 97% of the 343 quality-controlled *Porphyromonas* and outgroup genomes and was therefore included in phylogenomic tree construction. Of the 71 genes, 59 met this criterion and were retained for phylogeny.

**Table S6. GTDB classification of 343 quality-controlled *Porphyromonas* genomes.** Taxonomic assignments for the 343 quality-controlled *Porphyromonas* genomes based on the GTDB-Tk classify_wf workflow (v2.4.1, GTDB release r226). Columns include our internal genome identifiers and GTDB taxonomy from domain to species.

**Table S7. Metadata for 343 quality-controlled *Porphyromonas* genomes.** Columns list Genome ID, Ecological Source (human oral, human non-oral, or non-human oral taxon), Assigned Species, NCBI Organism Name, HOMD ID, GTDB taxonomy (Family, Genus, Species), host and isolation details (Host, Isolation Source, Type Material), and assembly information (RefSeq Category, Assembly Level, Sequencing Technology, Assembly Submitter). Genomes ordered by Ecological Source and taxonomy to highlight human oral taxa.

**Table S8. Type strains for human *Porphyromonas* species.** Selected *Porphyromonas* type strains for competitive mapping across human oral, stool, skin, and vaginal metagenomes from each human. Column A-C: species. Strain ID, Assembly Accession. Column D: Primary body site after mapping.

**Table S9. Dereplication clusters and representative genomes of 191 human oral *Porphyromonas* genomes.** The table lists 84 dereplication clusters from 191 quality-filtered *Porphyromonas* genomes at 98% ANI using a simple greedy algorithm. Columns include Cluster ID (1 to 84), Cluster Size (number of genomes), Representative Genome (used for competitive mapping), and Member Genomes (all genomes in the cluster). Representative genomes were selected preferentially from isolates over metagenome-assembled genomes (MAGs), based on type or reference strain status, genome quality (e.g., contiguity), presence in HOMD, and host/isolation source. Clusters are ordered by decreasing cluster size.

**Table S10. Pairwise ANI among 191 high-quality human oral *Porphyromonas* genomes.** Pairwise average nucleotide identity (ANI) values for 191 high-quality *Porphyromonas* genomes calculated using pyANI with the ANIb method are presented. Rows and columns are labeled by Genome ID, with identity values ranging from 0.00 to 1.00. This matrix was used to validate pangenomic groupings and to guide dereplication at 98% ANI. Corresponding ANI values are visualized as a heatmap in Fig. S1.

**Table S11. Summary mapping metadata for 1,242 oral metagenomes.** Metadata includes the oral site, total quality-filtered reads, total mapped reads, fraction of mapped reads, and subject gender.

**Table S12. Breadth of coverage for representative dereplicated *Porphyromonas* genomes across 1,242 oral metagenomes.** The table shows the breadth of coverage for each of the representative dereplicated genomes (rows, labeled by Genome ID) across 1,242 metagenomes from 9 healthy oral sites and 1 periodontitis-associated site (subgingival plaque). Breadth of coverage is defined as the fraction of nucleotides in a genome covered at least 1x, with values ranging from 0.00 to 1.00. Mapping was performed competitively against the dereplicated reference set, and coverage metrics were extracted from the resulting profiles.

**Table S13. Genome-level relative abundance across 1,242 oral metagenomes.** This table reports the relative abundance of each representative dereplicated genome across 1,242 oral metagenomes from 9 healthy and one disease-associated site. Genome-level relative abundance was calculated as the Q2-Q3 mean depth of coverage of a genome divided by the sum of Q2-Q3 depths of coverage across all genomes detected in the same sample. Rows correspond to genomes and columns to metagenomes.

**Table S14. Relative abundance of *Porphyromonas* genomic groups across oral sites.** The table contains the aggregated genome relative abundance into 10 genomic groups across the 1,242 oral metagenomes. Relative abundance for each genomic group was calculated as the sum of the relative abundance of its constituent genomes (data from Table S13). Rows correspond to genomic groups, and columns to oral metagenomes. Values represent the fraction of total genomic abundance contributed by each group in a sample, ranging from 0.0% to 100.0%.

**Table S15. Oral site prevalence of *Porphyromonas* genomic groups.** The table reports the percent of samples in which each *Porphyromonas* genomic group is detected across 9 healthy-associated and one disease-associated oral site. A genomic group was considered present in a sample if at least one of its constituent genomes reached a breadth of coverage ≥25% (data from Table S12). Rows correspond to taxa and columns to oral sites. Values are expressed as a percentage of samples in which each genomic group is detected per site, with the total number of samples per site shown in the column headers. To ease comparison with the relative abundance from Fig. 4, genomic groups and sites are in the same order.

**Table S16. Pairwise statistics for *Porphyromonas* genomic groups across oral sites.** The table reports contingency-associated metrics, abundance correlations, and dominance comparisons for pairs of genomic groups. Rows correspond to taxon pairs within a site. Columns include sample counts for each detection category, odds ratios, and Fisher’s exact test statistics, Spearman rank correlations among informative samples, and centered log-ratio (CLR) dominance comparisons for co-detected pairs. False discovery rate (FDR) corrected p-values are provided for multiple-testing adjustment.

**Table S17. Completeness of metabolic modules across 84 reference *Porphyromonas* genomes.** Percent of completeness of metabolic modules for each of the 84 reference genomes for modules where at least 75% of the necessary enzymes to complete the pathway are present. 43 of 188 modules are complete. Columns A and B include module ID and description; columns C-CH show percentages ranging from 75% to 100%.

**Table S18. Quality assessment of non-RefSeq *Porphyromonas pasteri* genomes.** The table summarizes genome quality metrics for 21 non-RefSeq *P. pasteri* genomes. Columns include GTDB Classification (v2.4.1; r226), Bacteria_71 Completion (≥70%), Bacteria_71 Redundancy (<10%), CheckM v1.2.3 Completeness (≥90%), CheckM v1.2.3 Contamination (<5%), CheckM2 v1.1.0 Completeness (≥90%), CheckM2 v1.1.0 Contamination (<5%), and Quality Status (Passed/Not Passed), indicating whether a genome met all thresholds. Seven out of 21 genomes passed all quality criteria.

**Table S19. Dereplication of RefSeq and non-RefSeq *Porphyromonas pasteri* genomes for site preference validation.** The table summarizes dereplication of all *P. pasteri* genomes used for taxon-specific competitive mapping. Dereplication was performed at 98% ANI across RefSeq and non-RefSeq genomes. Columns include Cluster ID, Cluster Size, Representative Genome, Genomes in Cluster, and NCBI Category (RefSeq/non-RefSeq). The seven non-RefSeq genomes formed their own clusters. Dereplication resulted in 24 *P. pasteri* clusters (7 non-RefSeq and 17 RefSeq).

Table S20. Pfam functional enrichment report in *Porphyromonas pasteri* genomes specialized to the tongue dorsum or supragingival plaque. Results of Pfam functional enrichment analysis in *P. pasteri* genomes are primarily distributed in tongue dorsum (TD) or supragingival plaque (SUPP). Columns include the source (Pfam), Enrichment Score, Unadjusted p-value, Adjusted q-value, Associated Groups (TD, SUPP, NA), Function Accession ID, Associated Gene Cluster IDs, counts of genomes with the function in each group (p_SUPP_clade, p_TD_clade, N_SUPP_clade, N_TD_clade), and validation flags (q-value <0.05, present in ≥50% of enriched group, present in <50% of non-enriched group, enriched with high ratio). Only one Pfam function was significantly enriched. See Methods “Functional enrichment analysis” for details.

Table S21. COG20 functional enrichment report in *Porphyromonas pasteri* genomes specialized to the tongue dorsum or supragingival plaque. COG20 functional enrichment results for *P. pasteri* gene clusters from TD and SUPP. Columns are as in Table S20. Only one COG20 **function was significantly enriched.**

Table S22. KOfam functional enrichment report in *Porphyromonas pasteri* genomes associated with tongue dorsum or supragingival plaque. KOfam functional enrichment results for *P. pasteri* gene clusters from TD and SUPP. Columns are as in Table S20. No functions were significantly enriched.

Table S23. CAZyme functional enrichment report in *Porphyromonas pasteri* genomes specialized to the tongue dorsum or supragingival plaque. CAZyme functional enrichment results for *P. pasteri* gene clusters from TD and SUPP. Columns are as in Table S20. No functions were significantly enriched.

Table S24. KEGG Module functional enrichment report in *Porphyromonas pasteri* genomes specialized to the tongue dorsum or supragingival plaque. KEGG Module functional enrichment results for *P. pasteri* gene clusters from tongue dorsum (TD) and supragingival plaque (SUPP). Columns are as in Table S20. No KEGG MODULES were significantly enriched.

**Table S25. Annotation of *P*. *gingivalis* mobile genomic island.** Gene call coordinates and functional annotations for 45 genes identified as part of a conjugative element in *P*. *gingivalis* ‘ATCC 33277’ along with flanking genes for reference. Annotations included: Pfam, NCBI COG20 (category, function, pathway), KEGG (KOfam, Module, BRITE, Class), CAZyme.

**Table S26. Nucleotide-level BLAST results for the mobile genomic island and flanking genes.** The table contains the top 3 BLASTn results for each gene using HOMD genomes as reference, ranked by identity (%), query coverage (%), and lowest e-value (columns C-F). Taxonomic information is provided for family, genus, and species level (column G-J). Human Microbe Taxon identifier is shown in column J.

**Table S27. Protein-level BLAST results for the mobile genomic island and flanking genes.** Genes were translated into amino acid sequences, and a protein-protein blast was performed. The table contains the top 3 BLASTp results for each gene using all annotated proteins in HOMD as reference, ranked by identity (%), query coverage (%), and lowest e-value (columns C-F). Taxonomic information is provided for family, genus, and species level (column G-J). Human Microbe Taxon identifier is shown in column J.

## Notes

### Competing Interest Statement

The authors have declared no competing interest.

## References

1. Costello EK, Lauber CL, Hamady M, Fierer N, Gordon JI, Knight R. 2009. Bacterial Community Variation in Human Body Habitats Across Space and Time. Science 326:1694–1697.

2. Dewhirst FE, Chen T, Izard J, Paster BJ, Tanner ACR, Yu W-H, Lakshmanan A, Wade WG. 2010. The Human Oral Microbiome. J Bacteriol 192:16.

3. The Human Microbiome Project Consortium. 2012. Structure, function and diversity of the healthy human microbiome. Nature 486:207–214.

4. Mark Welch JL, Dewhirst FE, Borisy GG. 2019. Biogeography of the Oral Microbiome: The Site-Specialist Hypothesis. Annu Rev Microbiol 73:335–358.

5. Socransky SS, Manganiello SD. 1971. The Oral Microbiota of Man From Birth to Senility. Journal of Periodontology 42:485–496.

6. Mark Welch JL, Rossetti BJ, Rieken CW, Dewhirst FE, Borisy GG. 2016. Biogeography of a human oral microbiome at the micron scale. Proc Natl Acad Sci USA 113.

7. Utter DR, Borisy GG, Eren AM, Cavanaugh CM, Mark Welch JL. 2020. Metapangenomics of the oral microbiome provides insights into habitat adaptation and cultivar diversity. Genome Biol 21:293.

8. Shaiber A, Willis AD, Delmont TO, Roux S, Chen L-X, Schmid AC, Yousef M, Watson AR, Lolans K, Esen ÖC, Lee STM, Downey N, Morrison HG, Dewhirst FE, Mark Welch JL, Eren AM. 2020. Functional and genetic markers of niche partitioning among enigmatic members of the human oral microbiome. Genome Biology 21:292.

9. McLean AR, Torres-Morales J, Dewhirst FE, Borisy GG, Mark Welch JL. 2022. Site-tropism of streptococci in the oral microbiome. Molecular Oral Microbiology 37:229–243.

10. Giacomini JJ, Torres-Morales J, Dewhirst FE, Borisy GG, Mark Welch JL. 2023. Site Specialization of Human Oral *Veillonella* Species. Microbiol Spectr 11:e04042–22.

11. Torres-Morales J, Mark Welch JL, Dewhirst FE, Borisy GG. 2023. Site-specialization of human oral *Gemella* species. Journal of Oral Microbiology 15:2225261.

12. Giacomini JJ, Torres-Morales J, Tang J, Dewhirst FE, Borisy GG, Mark Welch JL. 2024. Spatial ecology of *Haemophilus and Aggregatibacter* in the human oral cavity. Microbiol Spectr 12:e04017–23.

13. Giacomini JJ, Torres-Morales J, Dewhirst FE, Borisy GG, Mark Welch JL. 2025. Spatial ecology of the *Neisseriaceae* family in the human oral cavity. Microbiol Spectr 13.

14. Gibson FC, Genco CA. 2006. The Genus Porphyromonas, p. 428–454. In Dworkin, M, Falkow, S, Rosenberg, E, Schleifer, K-H, Stackebrandt, E (eds.), The Prokaryotes. Springer New York, New York, NY.

15. Paster BJ, Dewhirst FE, Olsen I, Fraser GJ. 1994. Phylogeny of Bacteroides, Prevotella, and Porphyromonas spp. and related bacteria. J Bacteriol 176:725–732.

16. Shah HN, Collins MD. 1988. Proposal for Reclassification of Bacteroides asaccharolyticus, Bacteroides gingivalis, and Bacteroides endodontalis in a New Genus, Porphyromonas. International Journal of Systematic Bacteriology 38:128–131.

17. Willems A, Collins MD. 1995. Reclassification of Oribaculum catoniae (Moore and Moore 1994) as Porphyromonas catoniae comb. nov. and Emendation of the Genus Porphyromonas. Int J Syst Bacteriol 45.

18. Sakamoto M, Li D, Shibata Y, Takeshita T, Yamashita Y, Ohkuma M. 2015. Porphyromonas pasteri sp. nov., isolated from human saliva. International Journal of Systematic and Evolutionary Microbiology 65:2511–2515.

19. Griffen AL, Becker MR, Lyons SR, Moeschberger ML, Leys EJ. 1998. Prevalence of *Porphyromonas gingivalis* and Periodontal Health Status. J Clin Microbiol 36:3239–3242.

20. Paster BJ, Boches SK, Galvin JL, Ericson RE, Lau CN, Levanos VA, Sahasrabudhe A, Dewhirst FE. 2001. Bacterial Diversity in Human Subgingival Plaque. J Bacteriol 183:3770–3783.

21. Abusleme L, Dupuy AK, Dutzan N, Silva N, Burleson JA, Strausbaugh LD, Gamonal J, Diaz PI. 2013. The subgingival microbiome in health and periodontitis and its relationship with community biomass and inflammation. ISME J 7:1016–1025.

22. Eren AM, Borisy GG, Huse SM, Mark Welch JL. 2014. Oligotyping analysis of the human oral microbiome. Proc Natl Acad Sci USA 111.

23. Guilloux C-A, Lamoureux C, Beauruelle C, Héry-Arnaud G. 2021. Porphyromonas: A neglected potential key genus in human microbiomes. Anaerobe 68:102230.

24. Galtier A, Warinner C, Velsko IM. 2026. Ancient species diversity and niche adaptation in Tannerella and Porphyromonas revealed through pangenomics. bioRxiv 10.64898/2026.02.09.704811.

25. Delmont TO, Eren AM. 2018. Linking pangenomes and metagenomes: the Prochlorococcus metapangenome. PeerJ 6:e4320–e4320.

26. Delmont TO, Kiefl E, Kilinc O, Esen OC, Uysal I, Rappe MS, Giovannoni S, Eren AM. 2019. Single-amino acid variants reveal evolutionary processes that shape the biogeography of a global SAR11 subclade. eLife 26.

27. Naito M, Hirakawa H, Yamashita A, Ohara N, Shoji M, Yukitake H, Nakayama K, Toh H, Yoshimura F, Kuhara S, Hattori M, Hayashi T, Nakayama K. 2008. Determination of the Genome Sequence of Porphyromonas gingivalis Strain ATCC 33277 and Genomic Comparison with Strain W83 Revealed Extensive Genome Rearrangements in P. gingivalis. DNA Research 15:215–225.

28. Naito M, Sato K, Shoji M, Yukitake H, Ogura Y, Hayashi T, Nakayama K. 2011. Characterization of the Porphyromonas gingivalis conjugative transposon CTnPg1: determination of the integration site and the genes essential for conjugal transfer. Microbiology 157:2022–2032.

29. Hurst R, Meader E, Gihawi A, Rallapalli G, Clark J, Kay GL, Webb M, Manley K, Curley H, Walker H, Kumar R, Schmidt K, Crossman L, Eeles RA, Wedge DC, Lynch AG, Massie CE, Yazbek-Hanna M, Rochester M, Mills RD, Mithen RF, Traka MH, Ball RY, O’Grady J, Brewer DS, Wain J, Cooper CS. 2022. Microbiomes of Urine and the Prostate Are Linked to Human Prostate Cancer Risk Groups. European Urology Oncology 5:412–419.

30. Ximénez-Fyvie LA, Haffajee AD, Socransky SS. 2000. Comparison of the microbiota of supra- and subgingival plaque in health and periodontitis. Journal of Clinical Periodontology 27:648–657.

31. Hajishengallis G, Lamont RJ. 2012. Beyond the red complex and into more complexity: the polymicrobial synergy and dysbiosis (PSD) model of periodontal disease etiology. Molecular Oral Microbiology 27:409–419.

32. Cohan FM, Koeppel AF. 2008. The Origins of Ecological Diversity in Prokaryotes. Current Biology 18:R1024–R1034.

33. Morris JJ, Lenski RE, Zinser ER. 2012. The Black Queen Hypothesis: Evolution of Dependencies through Adaptive Gene Loss. mBio 3:e00036–12.

34. Gogarten JP, Townsend JP. 2005. Horizontal gene transfer, genome innovation and evolution. Nat Rev Microbiol 3:679–687.

35. Renno AJ, Shields RC, McLellan LK. 2025. Bacterial evolution in the oral microbiome: the role of conjugative elements and horizontal gene transfer. J Bacteriol 207:e00066–25.

36. Eren AM, Esen ÖC, Quince C, Vineis JH, Morrison HG, Sogin ML, Delmont TO. 2015. Anvi’o: an advanced analysis and visualization platform for ‘omics data. PeerJ 3:e1319.

37. Eren AM, Kiefl E, Shaiber A, Veseli I, Miller SE, Schechter MS, Fink I, Pan JN, Yousef M, Fogarty EC, Trigodet F, Watson AR, Esen ÖC, Moore RM, Clayssen Q, Lee MD, Kivenson V, Graham ED, Merrill BD, Karkman A, Blankenberg D, Eppley JM, Sjödin A, Scott JJ, Vázquez-Campos X, McKay LJ, McDaniel EA, Stevens SLR, Anderson RE, Fuessel J, Fernandez-Guerra A, Maignien L, Delmont TO, Willis AD. 2021. Community-led, integrated, reproducible multi-omics with anvi’o. Nat Microbiol 6:3–6.

38. O’Leary NA, Cox E, Holmes JB, Anderson WR, Falk R, Hem V, Tsuchiya MTN, Schuler GD, Zhang X, Torcivia J, Ketter A, Breen L, Cothran J, Bajwa H, Tinne J, Meric PA, Hlavina W, Schneider VA. 2024. Exploring and retrieving sequence and metadata for species across the tree of life with NCBI Datasets. Sci Data 11.

39. Parks DH, Imelfort M, Skennerton CT, Hugenholtz P, Tyson GW. 2015. CheckM: assessing the quality of microbial genomes recovered from isolates, single cells, and metagenomes. Genome Res 25:1043–1055.

40. Chklovski A, Parks DH, Woodcroft BJ, Tyson GW. 2023. CheckM2: a rapid, scalable and accurate tool for assessing microbial genome quality using machine learning. Nature Methods 20:1203–1212.

41. Lee MD. 2019. GToTree: a user-friendly workflow for phylogenomics. Bioinformatics 35:4162–4164.

42. Edgar RC. 2004. MUSCLE: a multiple sequence alignment method with reduced time and space complexity. BMC Bioinformatics 5:113.

43. Capella-Gutierrez S, Silla-Martinez JM, Gabaldon T. 2009. trimAl: a tool for automated alignment trimming in large-scale phylogenetic analyses. Bioinformatics 25:1972–1973.

44. Nguyen L-T, Schmidt HA, von Haeseler A, Minh BQ. 2015. IQ-TREE: A Fast and Effective Stochastic Algorithm for Estimating Maximum-Likelihood Phylogenies. Molecular Biology and Evolution 32:268–274.

45. Minh BQ, Nguyen MAT, Von Haeseler A. 2013. Ultrafast Approximation for Phylogenetic Bootstrap. Molecular Biology and Evolution 30:1188–1195.

46. Yu G, Smith DK, Zhu H, Guan Y, Lam TT. 2017. GGTREE : an R package for visualization and annotation of phylogenetic trees with their covariates and other associated data. Methods Ecol Evol 8:28–36.

47. Parks DH, Chuvochina M, Rinke C, Mussig AJ, Chaumeil P-A, Hugenholtz P. 2022. GTDB: an ongoing census of bacterial and archaeal diversity through a phylogenetically consistent, rank normalized and complete genome-based taxonomy. Nucleic Acids Research 50:D785–D794.

48. Chaumeil P-A, Mussig AJ, Hugenholtz P, Parks DH. 2022. GTDB-Tk v2: memory friendly classification with the genome taxonomy database. Bioinformatics 38:5315–5316.

49. Hyatt D, Chen G-L, LoCascio PF, Land ML, Larimer FW, Hauser LJ. 2010. Prodigal: prokaryotic gene recognition and translation initiation site identification. BMC Bioinformatics 11:119.

50. Galperin MY, Wolf YI, Makarova KS, Vera Alvarez R, Landsman D, Koonin EV. 2021. COG database update: focus on microbial diversity, model organisms, and widespread pathogens. Nucleic acids research 49:D274–D281.

51. Camacho C, Coulouris G, Avagyan V, Ma N, Papadopoulos J, Bealer K, Madden TL. 2009. BLAST+: architecture and applications. BMC Bioinformatics 10:421.

52. Mistry J, Chuguransky S, Williams L, Qureshi M, Salazar GA, Sonnhammer EL, Tosatto SC, Paladin L, Raj S, Richardson LJ. 2021. Pfam: The protein families database in 2021. Nucleic acids research 49:D412–D419.

53. Kanehisa M, Goto S. 2000. KEGG: kyoto encyclopedia of genes and genomes. Nucleic acids research 28:27–30.

54. Kanehisa M, Sato Y, Morishima K. 2016. BlastKOALA and GhostKOALA: KEGG Tools for Functional Characterization of Genome and Metagenome Sequences. Journal of Molecular Biology 428:726–731.

55. Drula E, Garron M-L, Dogan S, Lombard V, Henrissat B, Terrapon N. 2022. The carbohydrate-active enzyme database: functions and literature. Nucleic Acids Research 50:D571–D577.

56. Chan PP, Lowe TM. 2019. tRNAscan-SE: Searching for tRNA Genes in Genomic Sequences, p. 1–14. In Kollmar, M (ed.), Gene Prediction. Springer New York, New York, NY.

57. van Dongen S, Abreu-Goodger C. 2012. Using MCL to Extract Clusters from Networks, p. 281–295. In van Helden, J, Toussaint, A, Thieffry, D (eds.), Bacterial Molecular Networks: Methods and Protocols. Springer New York, New York, NY.

58. Pritchard L, Glover RH, Humphris S, Elphinstone JG, Toth IK. 2016. Genomics and taxonomy in diagnostics for food security: soft-rotting enterobacterial plant pathogens. Anal Methods 8:12–24.

59. Califf KJ, Schwarzberg-Lipson K, Garg N, Gibbons SM, Caporaso JG, Slots J, Cohen C, Dorrestein PC, Kelley ST. 2017. Multi-omics Analysis of Periodontal Pocket Microbial Communities Pre- and Posttreatment. mSystems 2:e00016–17.

60. Minoche AE, Dohm JC, Himmelbauer H. 2011. Evaluation of genomic high-throughput sequencing data generated on Illumina HiSeq and Genome Analyzer systems. Genome Biol 12:R112.

61. Eren AM, Vineis JH, Morrison HG, Sogin ML. 2013. A Filtering Method to Generate High Quality Short Reads Using Illumina Paired-End Technology. PLoS ONE 8:e66643.

62. Langmead B, Salzberg SL. 2012. Fast gapped-read alignment with Bowtie 2. Nat Methods 9:357–359.

63. Li H, Handsaker B, Wysoker A, Fennell T, Ruan J, Homer N, Marth G, Abecasis G, Durbin R, 1000 Genome Project Data Processing Subgroup. 2009. The Sequence Alignment/Map format and SAMtools. Bioinformatics 25:2078–2079.

